# Screening of yeast display libraries of enzymatically-cyclized peptides to discover macrocyclic peptide ligands

**DOI:** 10.1101/2020.08.08.242537

**Authors:** John Bowen, John Schneible, Collin Labar, Stefano Menegatti, Balaji M. Rao

## Abstract

We present the construction and screening of yeast display libraries of cyclic peptides wherein site-selective enzymatic cyclization of linear peptides is achieved using bacterial transglutaminase. To this end, we developed two alternative routes, namely *(i)* yeast display of linear peptides followed by treatment with recombinant transglutaminase in solution; or *(ii)* intracellular co-expression of linear peptides and transglutaminase to achieve cyclization in the endoplasmic reticulum prior to yeast surface display. The cyclization yield was evaluated via orthogonal detection of epitope tags integrated in the yeast-displayed peptides by flow cytometry, and via comparative cleavage of cyclic *vs.* linear peptides by tobacco etch virus (TEV) protease. Subsequently, yeast display libraries of transglutaminase-cyclized peptides were screened to isolate binders to the N-terminal region of the Yes-Associated Protein (YAP) and its WW domains using magnetic selection and fluorescence activated cell sorting (FACS). The identified cyclic peptide cyclo[*E-*LYLAYPAH*-K*] featured a K_D_ of 1.67 µM for YAP and 0.84 µM for WW as well as high binding selectivity against albumin and lysozyme. These results demonstrate the usefulness of yeast surface display for screening transglutaminase-cyclized peptide libraries, and efficient identification of cyclic peptide ligands.

## Introduction

Cyclized peptide ligands feature superior bioactivity and biochemical stability compared to their linear counterparts.^1–4^ An extensive body of literature exists on cyclic peptides with antibody-like affinity for protein targets,^5,6^ the ability to permeate cells and deliver a therapeutic payload,^7,8^ or act as modulators of intracellular and extracellular protein-protein interactions.^9,10^ These studies emphasize the usefulness and the need for efficient methods to develop cyclic peptides with strong biorecognition activity. Several platform technologies have been developed for isolation of bioactive peptides by combinatorial library screening, such as mRNA-display,^11^ phage-display,^12^ yeast display,^13^ or screening of solid-phase synthetic libraries.^14^ However, peptide cyclization has been emphasized to a lesser extent,^15–19^ and has been predominantly achieved by the formation of disulfide bonds or nucleophilic substitution using bifunctional linkers on lysine and cysteine residues.^3,6,20,215,22,23^ More recently, enzymatic cyclization has been used to construct peptide macro-cycles on the surface of yeast^24^ and bacteria,^25^ or within bacteria.^26^ In one study, a library of lanthipeptides was cyclized using the ProcM enzyme, resulting in macrocyclic peptides displayed on the surface of yeast cells that were screened to identify ligands for αvβ3 integrin.^24^ In another study, peptides expressed on *E. coli* cells were combined with synthetic segments via intein trans-splicing and intramolecular oxime ligation.^25^ Another method proposed for intracellular synthesis of cyclic peptide libraries relies on split intein-mediated circular ligation of peptides and protein (SICLOPPS), and was employed to identify inhibitors to a variety of enzymes and protein-protein interactions.^27–29^

These enzymatic ligation strategies are gaining prominence owing to their excellent site- and chemo-selectivity, and high yields under mild conditions.^30–33^ As a result, new cyclization strategies based on specific peptide motifs are being developed: peptide macrocycles of up to 78 amino acids were produced by reacting proteins displaying an N-terminal glycine with the peptide motif LPXTG using Sortase;^34^ similarly, Butelase 1 has been employed to cyclize peptides ranging between 26 to greater than 200 residues;^35^ the S-adenosylmethionine enzyme AlbA has been used to catalyze the formation of a thioether bond for the synthesis of macrocyclic subtilosin A;^36^ finally, transglutaminase has been used in conjunction with orthogonal cyclization strategies to achieve poly-cyclization of peptides of varying lengths on the surface of phages.^37^

In this study, we present a method for constructing and screening yeast display libraries of transglutaminase-cyclized peptides to identify cyclic peptide ligands, using specific regions of the Yes-Associated Protein (YAP) as model targets. Transglutaminase catalyzes the formation of an amide (*i.e.*, peptide) bond between a glutamine (Q) residue located on the N-terminus of the variable region of the peptide and the C-terminal lysine (K) residue, resulting in a “head-to-side chain”-type peptide cyclization. In this study, the linear precursor peptides were displayed via the Aga1-Aga2 construct on the surface of yeast *Saccharomyces cerevisiae*.^13^ Specifically, we attempted two routes to achieve the cyclization of the linear peptide precursors with transglutaminase. The first one comprised the expression of the peptide-Aga fusion on yeast followed by the incubation of the cells with soluble recombinant transglutaminase. The second route relied on the co-expression of the linear peptides and transglutaminase by the same yeast cells with the aim of achieving peptide cyclization during intracellular trafficking through the endoplasmic reticulum and prior to display.^38^ The yield of transglutaminase-mediated cyclization was evaluated by comparing the detection levels of multiple epitope tags integrated in the peptide-Aga fusion via fluorescent immunolabeling and flow cytometry. Further confirmation of peptide cyclization was conducted by comparing the cleavage of transglutaminase-cyclized peptides by tobacco etch virus (TEV) protease against their linear precursors.

The resulting libraries were screened against the N-terminal region of the Yes-Associated Protein (YAP, AAs 1-291) and its WW domains (AAs 165-271) using magnetic selection and fluorescence associated cell sorting (FACS). YAP is a transcriptional regulator that plays a crucial role in cell proliferation and apoptosis.^39^ Ligands targeting YAP can serve as modulators of the Hippo signaling pathway, providing control over cellular mechanisms responsible for organ and tumor development and growth. The peptide cyclo[*E-*LYLAYPAH*-K*], identified by screening libraries of peptides cyclized via extracellular treatment with transglutaminase, was characterized via yeast display binding assay to determine the apparent affinity for the N-terminal domain of YAP (K_D_ ~ 1.67 μM) and its WW domains (K_D_ ~ 0.84 μM). Chromatographic assays were also conducted to evaluate binding affinity and selectivity on solid phase. Collectively, these results demonstrate the potential of this platform for the rapid *de novo* identification of cyclic peptides with protein biorecognition activity.

## Experimental

### Plasmids and yeast cell culture

The pCTCON vector containing the TRP selectable marker was used in conjugation with *Saccharomyces cerevisiae* strain EBY100. Specifically, the Frozen-EZ yeast transformation Kit II (Zymo Research) was used to transform plasmid DNA into chemically competent EBY100. Trp-deficient SDCAA and SGCAA media were used, respectively, for culturing and inducing cells harboring the pCTCON plasmid as previously described.^40^ During culture, yeast cells were grown in SDCAA media at 30°C while shaking at 250 RPM. To induce protein expression, yeast cells were transferred into SGCAA media at OD_600_ of 1 and incubated for 24 hrs or 48 hrs at 20°C under shaking at 250 rpm.

### Plasmid construction for yeast surface display of model peptides, the N-terminal region of YAP, and its WW domain

Plasmids were constructed using pCTCON as backbone vector to express the peptide sequences on the surface of yeast as Aga2 fusions. The pCTCON plasmid was digested at the NheI and BamHI sites. The pCTCON-YESS-pep construct affording the expression of the model peptide ALQSGSRGGGEQK was constructed by amplifying gene block 1 with Pf1 and Pr1. The pCTCON-YESS-pep was then used to incorporate the Gal10 transglutaminase enzyme. Gene block 2 was amplified with Pf2 and pr2. The resulting PCR product and the pTCTON-YESS-pep vector were digested with AgeI and KpnI, and ligated together to afford the co-expression of the Gal1 YESS peptide and Gal10 transglutaminase enzyme simultaneous in the pCTCON-YESS-Full-Length-TG-dual-display vector.

The plasmid pET22b(+) was utilized for recombinant expression of YAP and its WWdomain. Gene block 3 was amplified with Pf3 and Pr3, and inserted between the NdeI and XhoI sites of pET22b(+) resulting in the plasmid pET22b(+)-WW, which affords the expression of soluble WW domain (AAs 165-271). The plasmid pET22b(+) was also utilized to express the N-terminal region of YAP (AAs 1-291). Specifically, gene block 3 was amplified with Pf4 and Pr4 and inserted between the NdeI and XhoI sites of pET22b(+) resulting in the plasmid pET22b(+)-YAP, which affords the expression of soluble YAP.

All double-stranded gene fragments were purchased from Integrated DNA technologies (IDT, Coralville, IA). The primer oligonucleotides were acquired from IDT or Eton Biosciences (San Diego, CA). The gene fragment and primer sequences are listed in **Tables S1** and **S2**, while the randomized library oligonucleotides are in **Table S3**. Phusion Polymerase (Thermo Fisher Scientific, Waltham, MA) was used for PCR reactions following the manufacturer’s protocols. Restriction digests of plasmid backbones and PCR products were executed at 37°C for 2 hrs using a 5-times excess of each restriction enzyme. Digested plasmid backbones were incubated with Antarctic phosphatase (New England Biolabs, Ipswich, MA) for 1 hr at 37°C. Digested plasmids and PCR products were purified using a 9K series gel and PCR extraction kit (BioBasic, Markham, ON, Canada). Overnight ligations using T4 DNA ligase (Promega, Madison, WI) were performed with the digested plasmid backbones and PCR product inserts. Ligations were transformed into chemically competent Novablue *E. coli* cells. The cells were made chemically competent using Mix&Go! *E. coli* transformation buffers (Zymo Research, Irvine, CA). The GeneJET™ plasmid miniprep kit (Thermo Fisher Scientific) was used to harvest the plasmids from overnight *E. coli* cultures.

### Site directed mutagenesis of the pCTCON backbone

The pCTCON-YESS-Full-Length-TG-dual-display was initially modified via site-directed mutagenesis. The pCTCON-YESS-Pro-KR-Active-TG-dual-display vector was constructed via amplification of the pCTCON-YESS-Full-Length-TG-dual-display vector with Pf5 and Pr5, enabling the insertion of a 6 base pair oligonucleotide between the pro sequence and the active transglutaminase sequence. An aliquot of 50 ng of purified PCR product was treated with 1 μL of T4 polynucleotide kinase (New England Biolabs) and 1 μL of T4 DNA ligase for 1 hr at room temperature followed by incubation at 65°C for 20 min. DNA was then treated for 5 min at room temperature with 1 μL of DpnI (New England Biolabs) prior to transformation into electrocompetent Novablue *E. coli* cells. The pCTCON-YESS-Active-TG-dual-display construct was constructed from the pCTCON-YESS-Full-Length-TG-dual-display vector via amplification with Pf6 and Pr6 to remove the pro-TG sequence. A similar treatment using polynucleotide kinase, T4 DNA ligase, and DpnI enzyme treatment was executed to introduce a point mutation into the substrate sequence where the glutamine (Q) residue was replaced with an alanine (A) residue in order to remove one of the critical residues involved in the transglutaminase cyclization reaction. The dual display vectors wherein glutamine (Q) was replaced with alanine (A) were constructed by amplifying the pCTCON-YESS-Pro-KR-Active-TG-dual-display vector with Pf7and Pr7 prior to kinase, ligase, and DpnI treatment to yield vector pCTCON-YESS-Pro-KR-Active-TG-Q-to-A-SDM-dual-display. Also, pCTCON-YESS-Active-TG-dual-display vector was amplified with Pf7 and Pr7 prior to kinase, ligase, and DpnI treatment to yield vector pCTCON-YESS-Active-TG-Q-to-A-SDM-dual-display.

### Construction of combinatorial yeast display libraries of cyclic peptides via extracellular transglutaminase-mediated cyclization

Double stranded gene fragments were purchased from IDT, while the primer oligonucleotides were acquired from IDT or Eton Biosciences. The sequences of the randomized library oligonucleotides and primers are in **Tables S2** and **S3**. The yeast surface display pCTON vector was used as a backbone to create the combinatorial library for extracellular cyclization, as done in prior work.^40^ Briefly, a randomized DNA gene block (oligo 1 in **Table S3**) comprising 8 randomized NNK codons was amplified via PCR with primer Pf9 and Pr9. A series of 30 PCR reactions were performed in a volume of 50 μL comprising 1 U of Phusion HF DNA polymerase (Thermo Fisher Scientific), 1X Phusion Buffer, 0.2 mM deoxynucleotide triphosphate (dNTPs) (Thermo Fisher Scientific), 0.1 μM of the forward and reverse primers, 3% dimethyl sulfoxide (DMSO), and 10 ng of the template DNA. The PCR was performed using the following conditions: initial denaturation at 98°C for 2 min, followed by 30 cycles of denaturation at 98°C for 10 sec, annealing at 62°C for 20 sec, extension at 72°C for 5 sec, and a final extension at 72°C for 10 min. The PCR products were purified by conducting 2 phenol/chloroform/isoamyl alcohol extractions followed by 1 chloroform extraction. The extracted DNA was incubated overnight in two volumes of ethanol and 1/10 volume of potassium acetate at −20°C. The DNA was pelleted via centrifugation at 12,000x g for 20 min, washed with 70% ethanol, centrifuged again, and washed with 100% ethanol. The resulting DNA pellet was resuspended in DI water and stored at −20°C prior to library construction. To construct the plasmid backbone, gene block 4 was first amplified with Pf8 and Pr8. The pCTCON and the amplified PCR product of gene block 4 were digested with AgeI and XhoI, and ligated to generate vector pCTCON-TG-8mer-template, which was used as template backbone for the TG8mer library. The pCTCON-TG-8mer-template was digested with SalI and BamHI restriction enzymes at 37°C for 2 hrs. The digested plasmid backbones were incubated with Antarctic phosphatase (New England Biolabs) for 1 hr at 37°C. Subsequently, 12 μg of PCR product and 4 μg of linearized pCTCON-TG-8mer-template vector were transformed into competent *Saccharomyces cerevisiae* strain EBY100 for a total of four transformation reactions as done in prior work.^40,41^ The diversity of the pCTCON-TG-8mer resulting library was estimated to be ~ 2.6·10^7^. The resulting library of yeast cells were propagated in Trp-deficient SDCAA media at 30°C while shaking at 250 RPM. To induce protein expression, yeast cells were transferred into SGCAA media at OD_600_ of 1 and incubated for 24 hrs at 20°C under shaking at 250 rpm.

Lyophilized recombinant transglutaminase (Zedira, Darmstadt, Germany) was reconstituted to a concentration of 12.8 mg/mL in PBS, pH 6, and stored as frozen aliquots at −20°C. Yeast cells displaying the library of linear peptide precursors were spun down and washed with PBS, pH 6. An aliquot of 10^8^ cells were resuspended in 500 μL of PBS, pH 6.0, to which the transglutaminase solution was added to achieve a final concentration of 1 μM. Cells were incubated at 37°C overnight under mild agitation. The cells were washed 3x with 0.1% BSA in PBS and analyzed via flow cytometry single cell analysis to evaluate the yield of peptide cyclization or screened against WW domain or YAP protein to identify peptide binders.

### Construction of combinatorial yeast display libraries of cyclic peptides via intracellular transglutaminase-mediated cyclization

The library for intracellular cyclization was constructed using a randomized DNA gene block (oligo 2 in **Table S3**) comprising 7 randomized NNK codons and amplified via PCR using primers Pf10 and Pr10. The PCR amplification was performed using the following conditions: initial denaturation at 98°C for 2 min, followed by 30 cycles of denaturation at 98°C for 10 sec, annealing at 62°C for 20 sec, extension at 72°C for 10 sec, and a final extension at 72°C for 10 min. The PCR products were purified via phenol/chloroform/isoamyl alcohol extractions and ethanol precipitation, as described above. The vector pCTCON-YESS-Active-TG-dual-display construct was digested with NheI and XhoI restriction enzymes at 37°C for 2 hrs. Digested plasmid backbones were incubated with Antarctic phosphatase (New England Biolabs) for 1 hr at 37°C. Subsequently, 12 μg of PCR product and 4 μg of linearized pCTCON-YESS-Active-TG-dual-display vector were transformed into competent *Saccharomyces cerevisiae* strain EBY100 as previously described.^40,41^ The diversity of the resulting pCTCON-TG-intracellular-7mer library was estimated to be ~ 1.6·10^7^. The resulting library of yeast cells were propagated in Trp-deficient SDCAA media at 30°C while shaking at 250 RPM. To induce protein expression, yeast cells were transferred into SGCAA media at OD_600_ of 1 and incubated for 48 hrs at 20°C under shaking at 250 rpm.

### Expression, purification, and biotinylation of recombinant N-terminal YAP, YAP-WW domain, and TEV

Plasmid pET22b(+)-WW was transformed into chemically competent Rosetta *E. coli* cells. A frozen aliquot of transformed Rosetta *E. coli* was grown in LB media (10 g/L tryptone, 5 g/L yeast extract, 10 g/L NaCl) overnight at 37°C. The cells were then inoculated in 2XYT media (16 g/L tryptone, 10 g/L yeast extract, and 5 g/L NaCl) and allowed to propagate at 37°C for ~5 hrs until an optical density of ~1.0 was obtained. Protein expression was induced by adding IPTG to a total concentration of 0.5 mM and was allowed to proceed for 20 hrs at 20°C. An identical strategy was executed on plasmids pET22b(+)-YAP and pRK793 (#8827 from Addgene) for the expression of YAP and tobacco etch virus (TEV) protease.

The N-terminal YAP, WW-YAP, and TEV proteins were purified by immobilized metal affinity chromatography (IMAC). The cells were initially pelleted by centrifugation at 3500xg for 12 min, and resuspended in 35 mL of resuspension buffer (20 mM HEPES, 150 mM NaCl, pH 7.8, 0.2 mM phenylmethylsulfonyl). The cell slurry was sonicated and subsequently pelleted via centrifugation at 15,000xg for 22 min. The supernatant was then syringe filtered using a 0.45 μm PVDF filter (Genesee Scientific, San Diego, CA). The filtered cell lysate was loaded onto a 5 mL Nuvia IMAC column (Bio-Rad, Hercules, CA) at 2 mL/min, washed with 50 mL of Buffer C-IMAC (20 mM HEPES, 800 mM NaCl, pH 7.8) at 5 mL/min, flushed with 50 mL of Buffer A-IMAC (20 mM HEPES, 137 mM NaCl, pH 7.8 at 5 mL/min), and eluted with a 50 mL linear gradient of 0-100% Buffer B-IMAC (20 mM HEPES, 137 mM NaCl, 500 mM Imidazole, pH 7.8) at 5 mL/min. The chromatographic fractions were analyzed for protein content by UV spectrophotometry at 280 nm, and the fractions containing the protein were further analyzed via SDS-PAGE. A volume of 4.5 μL of protein fraction was combined with 1.5 μL of 4x LDS sample buffer (Invitrogen), loaded into a 4-12% Bis-Tris NuPAGE protein gel (Invitrogen) with MES SDS running buffer, and run at 200 V for ~40 min. The gel was stained with Imperial protein stain (Thermo Fisher), washed 3x with water, and allowed to de-stain overnight in water under mild agitation. The gel was analyzed, and selected chromatographic fractions were pooled and dialyzed into 20 mM HEPES, 150 mM NaCl, pH 7.8 (HEP) using SnakeSkin™ Dialysis Tubing (MWCO 3.5 kDa, Thermo Fisher Scientific) prior to biotinylation. Proteins were biotinylated using a EZ-Link Sulfo-NHS-LC-Biotinylation kit (Thermo Fisher Scientific) following the manufacturer’s protocol. Excess biotin was removed via dialysis into 20 mM HEPES, 150 mM NaCl, pH 7.8 (HEP) using SnakeSkin™ Dialysis Tubing (MWCO 3.5 kDa, Thermo Fisher Scientific). The protein was flash-frozen in liquid nitrogen and stored at −80°C in HEP plus 10% glycerol for long-term storage.

### Evaluation of extracellular and intracellular transglutaminase-mediated peptide cyclization via flow cytometry

Yeast cells displaying cyclic peptides were grown in SDCAA media for 24 hrs, collected, and induced at an OD of ~1.0 in SGCAA for 24, 48, 72, or 96 hrs to determine the effect of induction time on cyclization efficiency. After induction, the cells were collected and labeled with a 1:100 dilution of the chicken anti-c-myc antibody (Invitrogen), rabbit anti-HA antibody (Thermo Fisher Scientific), and anti His-488 (Qiagen) or anti His-647 (Qiagen) conjugate; all antibodies were diluted in 0.1% w/v BSA in PBS, and incubated with the cells for 15 min at 4°C. Following incubation, the cells were washed, and secondary labeling was performed with goat anti-chicken 488, goat anti-chicken 633, donkey anti-rabbit 488, donkey anti-rabbit 633, or donkey anti-mouse 488, or donkey anti-mouse 633 (Thermo Fisher Scientific) at a 1:250 dilution for 15 min at 4°C. Cells were analyzed via fluorescent flow cytometry single cell analysis using a Biotec MACsQuant VYB cytometer (Miltenyi, Bergisch Gladbach, Germany). Labeling with secondary antibodies conjugated to a 488 fluorophore were imaged at the excitation wavelength of 488 nm and emission wavelength of 525/50 nm. Labeling with secondary antibodies conjugated to 633 or 647 fluorophores were imaged at the excitation wavelength of 561 nm and emission wavelength of 661/20 nm. At least 50,000 events were recorded for each sample. The mean fluorescence intensity (MFI) of each population was obtained using FlowJo flow cytometry software.

Yeast cells displaying cyclic peptides were spun down and washed 3x with 150 mM NaCl in PBS, pH 8. An aliquot of 10^8^ cells were resuspended in PBS, pH 8, to which recombinant TEV was added to achieve a final concentration of 3 μM. Cells were incubated overnight at 37°C under mild agitation. The cells were then washed 3x with 0.1% BSA in PBS and analyzed via flow cytometry as described above.

### Screening of yeast display libraries of transglutaminase-cyclized peptides via magnetic selection and FACS to identify affinity peptide ligands

The yeast display libraries of cyclic peptides produced by transglutaminase-mediated extracellular cyclization were subjected to four rounds of screenings to identify ligands to the WW domain and the N-terminal YAP. The first two rounds of screening were conducted by implementing magnetic selection. In round 1, 10^9^ library cells were incubated with the target protein immobilized onto magnetic beads. These were prepared by incubating 17 μg of biotinylated N-terminal YAP or WW-YAP with 25 μL of biotin binder Dynabeads (Invitrogen, Carlsbad, CA) for 2 hrs at room temperature; following conjugation, the beads were separated using a magnet and washed 3x with 0.1% BSA in PBS. In round 2, 10^8^ library cells were incubated with the same amount of target protein and bead volume. In both rounds, the pCTCON-TG-8mer library was initially induced for 24 hours whereas the pCTCON-TG-intracellular-7mer library was induced for 48 hour and washed with 0.1% BSA in PBS prior to selection, and then incubated with the protein-coated beads in 0.1% BSA in PBS for 1 hr at room temperature. Following incubation, the cells bound to the beads were isolated using a magnet, transferred into 5 mL of SDCAA media and allowed to propagate at 30°C for 2 days. Subsequently, 2 rounds of fluorescence activated cell sorting (FACS) were performed on a FACSAria II (Becton Dickinson, Franklin Lakes, NJ) cell sorter. Specifically, 10^7^ library cells were labeled with 1 μM YAP or WW-YAP protein for 1 hour at 4°C. Subsequently, cells were washed with 0.1% BSA in PBS, and resuspended in 1:250 dilution of Streptavidin-phycoerythrin (Invitrogen) for 15 minutes at 4°C. Cells were washed again with 0.1% BSA in PBS, and resuspended in 1 mL prior to sorting. Individual cells were selected based on their increased fluorescence intensity compared to a population of library cells that were not incubated with WW or YAP protein. During the second round of screening, cells were incubated with 100 nM of WW or YAP protein, and the cell collection gating stringency was increased by only collecting cells with a higher fluorescence intensity compared to the previous round. Following screening, the yeast cells were plated at varying dilutions on SDCAA agar plates and grown at 30°C for 2 days. Individual clones were selected, transferred into 5 mL of SDCAA liquid media and grown for 2 days. The yeast cell plasmid DNA was then extracted via a Zymoprep yeast miniprep kit (Zymo Research), propagated into electrocompetent Novablue *E. coli* cells, extracted using a GeneJET™ plasmid miniprep kit (Thermo Fisher Scientific), and finally sequenced.

### Yeast surface display isotherms of cyclo[E-LYLAYPAH-K] for N-terminal YAP and WW-YAP

Yeast cells displaying cyclo[*E-*LYLAYPAH*-K*] were prepared and incubated with varying concentrations of biotinylated WW-YAP or N-terminal YAP for 1 hr at 4°C. After washing with 0.1% BSA in PBS, the cells were labeled with a 1:250 dilution of R-Phycoerythrin-labeled Streptavidin. Flow cytometry analysis was executed using an excitation wavelength of 561 nm and an emission filter of 586/15 nm. The on-yeast binding measurements were fit to **Equation 1** via non-linear least squares regression over three independent replicates.

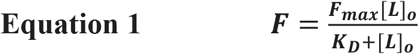

Where F is the mean fluorescence intensity, [L]_o_ is the concentration of the soluble protein used for the primary incubation, and F_max_ is the maximum fluorescence intensity when surface-saturation is obtained (*note:* the fluorescence signal was observed to decrease at high values of [L]_o_ (hook effect), as previously observed in yeast surface titrations;^42^ accordingly, in each dataset the highest value of fluorescence was set as F_max_ and the adsorption data obtained at higher protein concentration were excluded from the fittings).

### Synthesis of peptides cyclo[E-LYLAYPAH-K] and RYSPPPPYSSHS (PTCH) on Toyopearl resin

The peptides were synthesized on Toyopearl AF-Amino-650 M resin (0.2 meq/mL, Tosoh, Tokyo, Japan) using a Alstra Initator+ automated synthesizer (Biotage, Uppsala, Sweden) via standard solid-phase Fmoc/tBu chemistry.^43,44^ Prior to the coupling the first amino acid, the resin was swollen in DMF at 60 °C for 30 min. The linear precursor ELYLAYPAHK was synthesized using the sequence of protected amino acids Fmoc-Lys(Mtt)-OH, Fmoc-His(Trt)-OH, Fmoc-Ala-OH, Fmoc-Pro-OH, Fmoc-Tyr(OtBu)-OH, Fmoc-Ala-OH, Fmoc-Leu-OH, Fmoc-Tyr(OtBu)-OH, Fmoc-Leu-OH, and Fmoc-Glu(OAll)-OH. The PTCH peptide RYSPPPPYSSHS was synthesized using the sequence of Fmoc-Ser(OtBu)-OH, Fmoc-His(Trt)-OH, 2x Fmoc-Ser(OtBu)-OH, Fmoc-Tyr(OtBu)-OH, 4x Fmoc-Pro-OH, Fmoc-Ser(OtBu)-OH, Fmoc-Tyr(OtBu)-OH, and Fmoc-Arg(Pbf)-OH. All amino acids couplings were conducted with 5 equivalents (5 eq. compared to the functional density of Toyopearl AF-Amino-650 M resin) of Fmoc-protected amino acid (at the concentration of 0.5 M in anhydrous DMF), 4.95 eq. 2-(6-Chloro-1H-benzotriazole-1-yl)-1,1,3,3-tetramethylaminium hexafluorophosphate (HCTU, 0.5 M in DMF) and 10 eq. diisopropylethylamine (DIPEA, 2M in NMP) for 5 min at 75°C. Successful completion of every amino acid couplings was monitored via Kaiser’s ninhydrin assay.^45^ Fmoc deprotection was achieved by incubating the resin twice in 20% piperidine in DMF (v/v) at RT for 10 min. All Fmoc-protected amino acids, HCTU, piperidine, and anhydrous solvents were from Chem-Impex International, Inc. (Wood Dale, IL). The cyclization of QLYLAYPAHK was performed following a method developed in prior work.^46^ Briefly, the allyl ester (OAll) protection was removed from N-terminal Glu(OAll) using Tetrakis(triphenylphosphine)palladium(0) in DCM, while the methyltrityl (Mtt) protection was removed from C-terminal Lys(Mtt) in 2% trifluoroacetic acid (TFA) and 5% triisopropylsilane (TIPS) in DCM.^47^ The conjugation of the free carboxyl group of Glu and amino group of Lys via amide bond was performed by incubating the resin twice with 1 eq. HCTU (0.5 M in DMF) and 2 eq. DIPEA (2M in NMP) for 20 min at 75°C. All reagents utilized for site-selective deprotection and peptide cyclization were acquired from Millipore Sigma (Saint Louis, MO). Both peptides were rinsed copiously with DCM, and deprotected by acidolysis using a 90/5/3/2 TFA/Thioanisole/EDT/Anisole cocktail at 10 mL per gram of resin for 2 hrs at RT. The resin were then washed copiously with DCM, DMF, DCM, dried under nitrogen, and stored at −20 °C.

### Evaluation of binding capacity, affinity, and selectivity of resin-bound peptides to N-terminal YAP and WW-YAP

Aliquots of 1 mg of cyclo[*E-*LYLAYPAH*-K*]-Toyopearl resin and RYSPPPPYSSHS-Toyopearl resin were washed 3x with PBS, pH 7.4, and incubated with either N-terminal YAP or WW-YAP at concentrations varying from 20 μg/mL to 3 mg/mL in a volume of 400 μL for 1 hr at room temperature. After removing the resin by centrifugation, the supernatant was analyzed by spectrophotometry at 280 nm to quantify the concentration of unbound protein. The amount of protein bound to the peptide-Toyopearl resin was calculated by mass balance. The adsorption data were fit to a Langmuir isotherm (**Equation 2**):

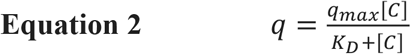

Wherein *q* is the amount of protein adsorbed on the peptide-Toyopearl resin (mg protein/mL resin), *[C]* is the unbound concentration of protein at equilibrium (mg protein/mL solution), *q_max_* is the maximum protein binding capacity (mg protein/mL resin), and *K_D_* is the dissociation constant (affinity, mg protein/mL solution).

The binding specificity of cyclo[*E-*LYLAYPAH*-K*] for WW-YAP was then evaluated. Bovine serum albumin (BSA) and lysozyme (LYZ) (Fisher Scientific) were dissolved in 20 mM HEPES, 150 mM NaCl, pH 7.8 and serially diluted to multiple concentrations ranging from 13 μg/mL to 2 mg/mL. An aliquot of 1 mg of cyclo[*E-*LYLAYPAH*-K*]-Toyopearl resin was washed in PBS, pH 7.4, and incubated in 400 μL of either BSA or LYZ solution for 1 hr at room temperature. After removing the resin by centrifugation, the supernatant was analyzed by spectrophotometry at 280 nm to quantify the concentration of unbound protein. The amounts of protein bound to the peptide-Toyopearl resin (q_BSA_ and q_LYZ_) were calculated by mass balance and compared to the amount of N-terminal YAP or WW-YAP that would be adsorbed at the same value of equilibrium protein concentration *[C]*.

## Results and Discussion

### Evaluation of extracellular transglutaminase-mediated peptide cyclization

Transglutaminase catalyzes the formation of an amide (*i.e.*, peptide) bond between the ε-amino group of a C-terminal lysine and the γ-carbamoyl side chain group of an N-terminal glutamine (**Figure 1A**), and has been widely utilized for protein modification, and peptide conjugation and cyclization.^48^ In the context of cyclization of short peptides, transglutaminase has been utilized either alone or in combination with orthogonal cyclization agents to construct cyclic and polycyclic constructs comprising 6 – 12 residues.^37,49^ Considerable efforts have been dedicated to the engineering of the enzyme,^50^ the identification of its recognition sequence, and the optimization of the reaction conditions to increase the yield of ligation;^37,51^ in particular, it has been recognized that appending the N-terminal dipeptide Ala-Leu significantly increases the cyclization yield.^37^

**Figure 1.**
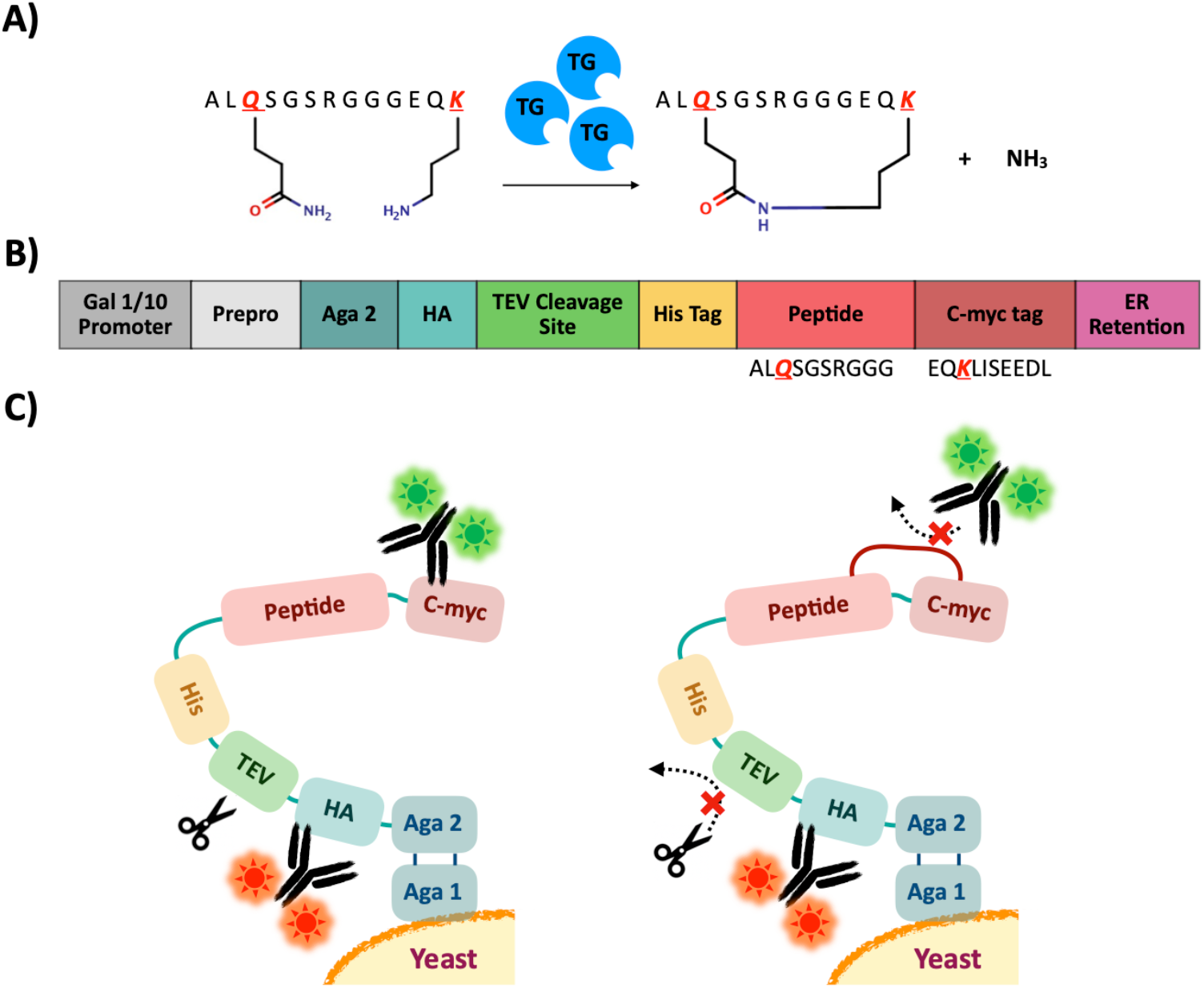
Transglutaminase-mediated cyclization of model peptides displayed on the surface of yeast cells and evaluation of peptide cyclization efficiency. **(A)** Formation of an amide bond between the ε-amino group of a C-terminal lysine and the γ-carbamoyl side chain group of an N-terminal glutamine by transglutaminase; **(B)** Design of the pCTcon2 plasmid vector for expressing the model peptide sequence ALQSGS-RGGGKS as contiguous to the c-myc epitope sequence EQKLISEEDL; **(C)** Scheme of the peptide construct and the impact of transglutaminase-mediate peptide cyclization on the detection of the c-myc and HA tags and the efficiency of cleavage by TEV protease.

In this study, we referred to the sequence ALQSGSRGGGKS as linear precursor to evaluate the efficiency of cyclization on the surface of yeast cells; this peptide, in fact, served as model substrate in prior work on transglutaminase-mediated cyclization.^37^ Specifically, we designed a plasmid construct to express the lead sequence ALQSGSRGGG in an Aga1-Aga2 display system on the surface of *Saccharomyces cerevisiae* cells. Together with the model peptide, the Aga2 fusion construct comprises the c-myc and HA detection tags, and a tobacco etch virus (TEV) protease signal recognition sequence (**Figure 1B**). Notably, the c-myc epitope EQKLISEEDL provides the lysine residue (K) that is enzymatically ligated with the glutamine (Q) of the lead sequence ALQSGSRGGG, resulting in peptide cyclization. We hypothesized that the transglutaminase-mediated tethering of the lead sequence and the c-myc epitope hinders – or completely prevents – the binding of c-myc antibodies (**Figure 1C**). Accordingly, the quantification of the differential binding of fluorescently labeled c-myc antibodies to cells displaying either linear or transglutaminase-cyclized peptides provides a measure of the cyclization yield (*note:* to ensure the site-selectivity of the cyclization, we eliminated the lysine and serine residues from the C-terminus of the model peptide sequence).

Upon inducing the expression of the linear peptide construct on the yeast surface, the cells were either treated with soluble recombinant transglutaminase or plain buffer (negative control). The expression of the peptide was confirmed by detecting the HA and c-myc tags via flow cytometry (**Figure 2A, D**). While the HA signal did not differ between transglutaminase-treated and untreated cells, a notable decrease in detection c-myc signal was found with the cells incubated with transglutaminase. This is consistent with our hypothesis of decreased antibody binding on c-myc tag upon enzymatic crosslinking to the model sequence.

**Figure 2.**
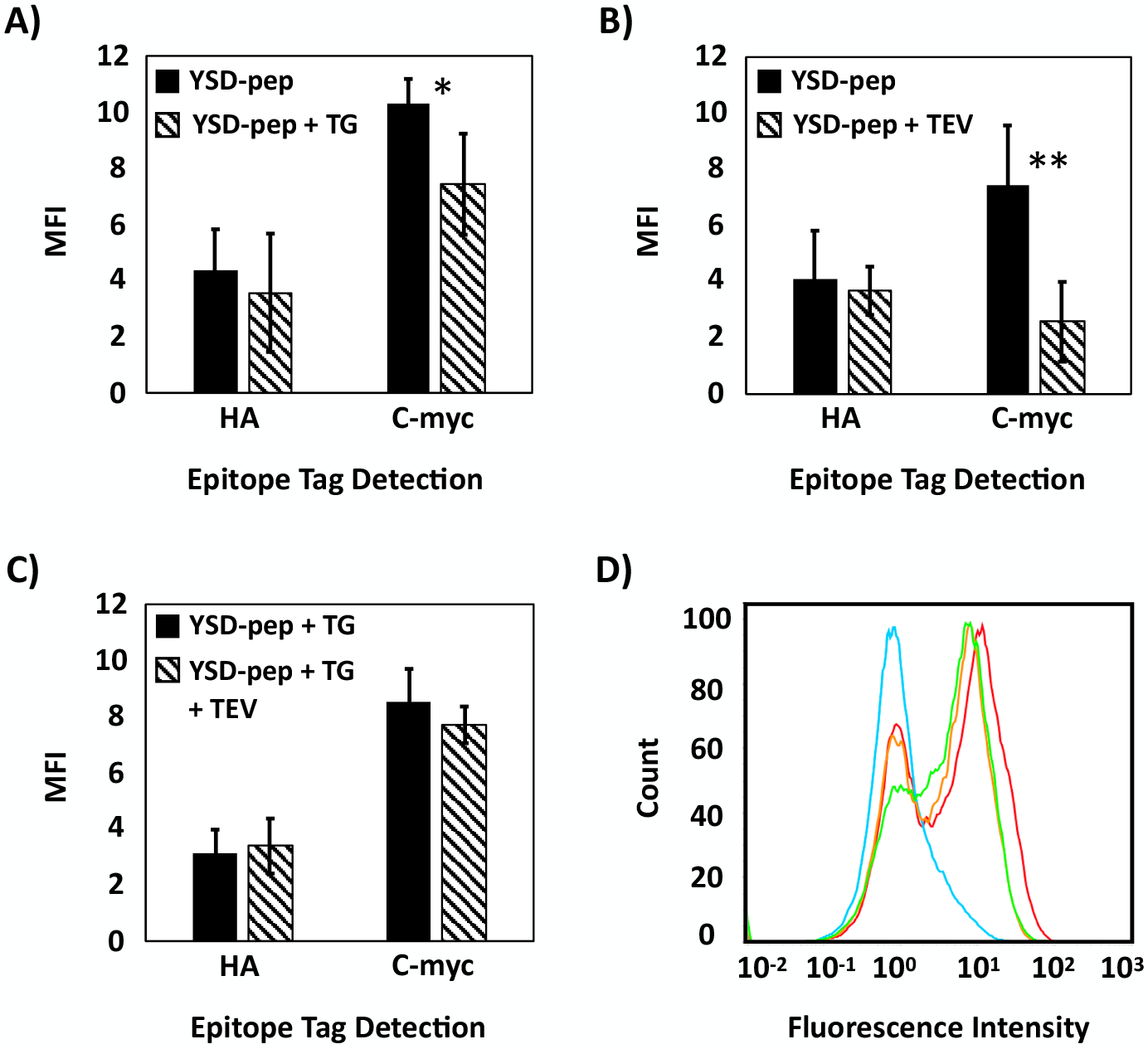
Evaluation of the yield of transglutaminase-mediated cyclization of yeast displayed peptides via immunofluorescence labeling and flow cytometry. **(A)** Mean fluorescence intensity (MFI) of HA and c-myc tag levels detected on yeast surface display peptides before (YSD-pep) and after (YSD-pep + TG) incubation of the yeast cells with transglutaminase (TG); **(B)** MFI of HA and c-myc tag levels detected on yeast surface display peptides before (YSD-pep) and after (YSD-pep + TEV) incubation of the yeast cells with TEV protease (no incubation with TG); **(C)** MFI of HA and c-myc tag levels detected on yeast surface display peptides incubated with TG before (YSD-pep + TG) and after (YSD-pep + TG + TEV) incubation of the yeast cells with TEV protease; **(D)** Representative flow cytometry histograms of c-myc epitope levels detected on yeast cells displaying the linear peptide before (red) and after (orange) incubation with TG, with no incubation with TG and after treatment with TEV protease (cyan), and after incubation with TG followed by treatment with TEV protease (green). The error bars correspond to the standard error of the mean from three independent replicates. A two-tailed paired t-test was performed; * indicates p<0.1 and ** indicates p<0.05.

We further evaluated the cyclization yield by assessing the extent of cleavage of linear *vs.* transglutaminase-treated peptide constructs from yeast cells by TEV protease. This cysteine protease recognizes the sequence ENLYFQ-G/S and cleaves it at the Q and G/S residues.^38^ We hypothesize that peptide cyclization would inhibit the cleavage of the TEV recognition sequence framed between the His-tag and the HA tag. Accordingly, yeast cells expressing the cyclized peptides or the linear precursor were treated with TEV protease, and the level of the HA and c-myc tags were quantified via fluorescent flow cytometry. No variation in HA signal upon treatment with TEV protease was observed in cells displaying either the linear or transglutaminase-cyclized peptide (**Figure 2B, C**); this was expected, given that the TEV recognition sequence is in down-stream to the HA tag (**Figure 1C**). However, a notable difference in residual c-myc level was observed between the cells displaying the linear *vs.* transglutaminase-cyclized peptides, wherein the former cell population showed a significant loss of fluorescence (**Figure 2B, D**) while the latter only presented a minor decrease (**Figure 2C, D**). This is coherent with our hypothesis that the TEV recognition site is significantly sterically hindered upon peptide cyclization or is in itself involved in the cyclization through its glutamine residue (*note:* the latter is unlikely due to the lack of Ala-Leu leader sequence required for transglutaminase reaction in the TEV recognition site). Collectively, these results suggest that extracellular treatment with transglutaminase results in the cyclization of the linear peptide precursor expressed as an Aga fusion on the surface of the yeast cells.

### Design, construction and screening of a yeast display library of cyclic peptides via extracellular transglutaminase-mediated cyclization

We implemented the extracellular peptide cyclization method demonstrated above to the construction of a yeast display library of cyclic 8-mer peptides using the randomized sequence WALQX_1_X_2_-X_3_-X_4_-X_5_-X_6_-X_7_-X_8_KS as linear precursor (**Figure 3A**, *note:* because a C-terminal lysine (K) was incorporated in the library sequence, the c-myc tag was omitted from the construct). The cyclization yield of the library peptides was evaluated via flow cytometry upon fluorescent labeling of the HA tag and His-tag, with and without treatment with TEV protease. As anticipated, no difference in HA tag levels was observed upon treatment with transglutaminase (**Figure 3B**). Exposure to TEV protease resulted in a notable decrease in HA tag levels for yeast display library of linear peptides (**Figure 3C**), whereas no significant variation was observed with the yeast display library of cyclic peptides (**Figure 3D**). These results suggest that cyclization of the peptide yeast library was effectively achieved with transglutaminase.

**Figure 3.**
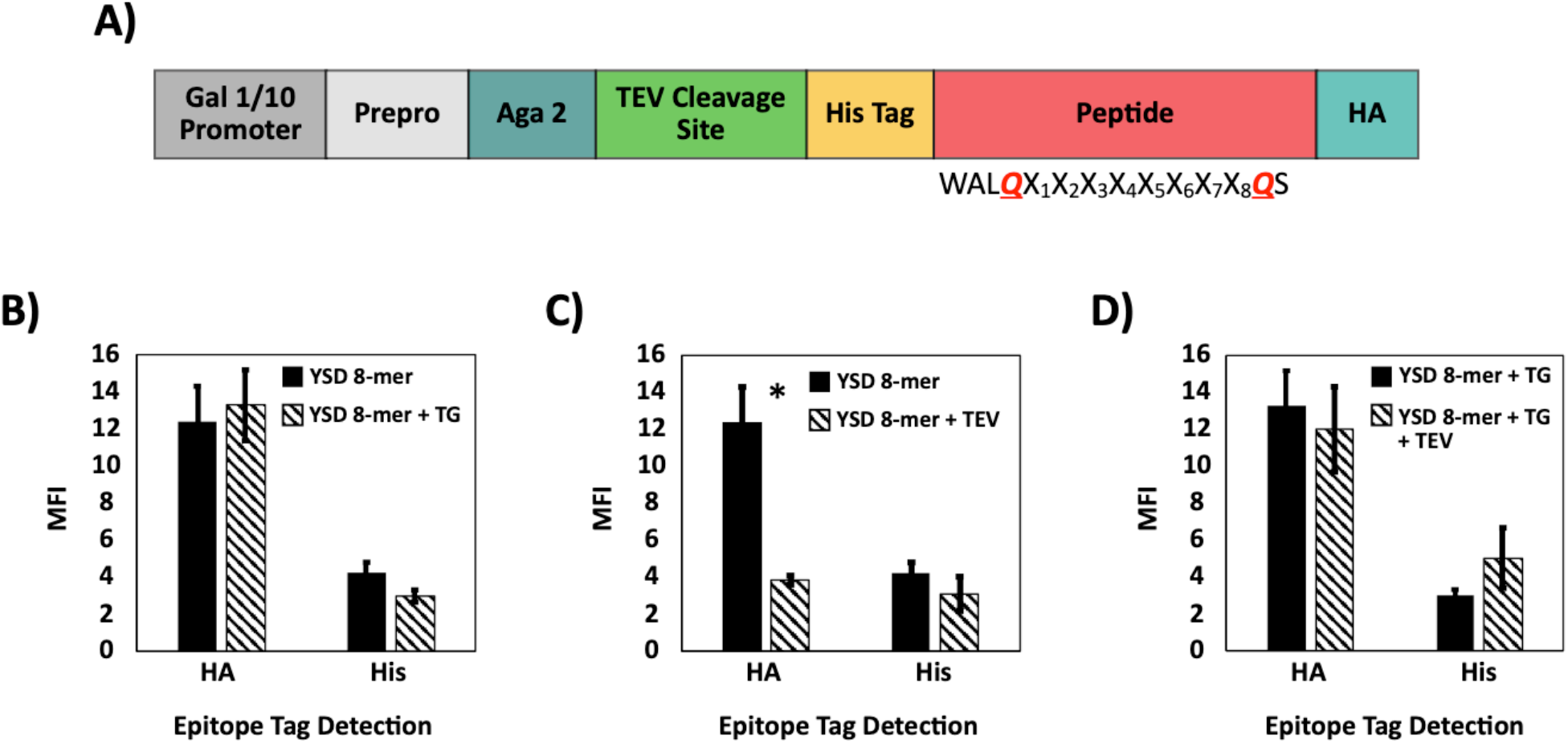
**(A)** Design of a pCTcon2 vector-based plasmid to express a combinatorial library of 8-mer peptides using the Aga2 display system on the surface of yeast cells; **(B)** MFI of HA and His tag levels detected on yeast surface display library of 8-mer peptides before (YSD 8-mer) and after (YSD 8-mer + TG) incubation of the yeast cells with transglutaminase (TG); **(C)** MFI of HA and His tag levels detected on a yeast surface display library of 8-mer peptides before (YSD 8-mer) and after (YSD 8-mer + TEV) incubation of the yeast cells with TEV protease (no incubation with TG); **(D)** MFI of HA and His tag levels detected on a yeast surface display library of 8-mer peptides incubated with TG before (YSD 8-mer + TG) and after (YSD 8-mer + TG + TEV) incubation of the yeast cells with TEV protease. Error bars correspond to the standard error of the mean from three independent replicates. A two-tailed paired t-test was performed. * p<0.05.

We then screened the library against the N-terminal region (AAs 1-291) of Yes-Associated Protein 65 (N-YAP) and its WW domains (WW-YAP). To ensure the selection of peptide ligands with true affinity for the target proteins, two rounds of screening via magnetic cell sorting followed by two rounds of screening via fluorescence activated cell sorting (FACS) were performed; the stringency of screening was increased during FACS by lowering the concentration of the target protein from 1 μM to 100 nM. The screening against each target returned 10 yeast clones from which the plasmid DNA was extracted, transformed into *E. coli* cells, and sequenced. Notably, 9 out of 10 clones obtained from screening against N-YAP and all the 10 clones obtained from screening against WW-YAP returned the peptide sequence QLYLAYPAHK. This corresponding to the cyclic peptide construct cyclo[*E*-LYLAYPAH-*K*], since the transglutaminase-mediated transamidation of the carbamoylethyl side chain group of glutamine (Q) with the ε-amine group of lysine (K) is equivalent to the formal amidation of the carboxyethyl side chain group of a glutamic acid (E) by the ε-amine group of lysine (K). This sequence features a defined amphiphilic character (*i.e.*, a balance of hydrophobic and polar residues) and, most notably, a combination of proline and tyrosine residues that is consistent with the PPxY motif characteristic of WW-binding peptides.

### Binding affinity and specificity of cyclo[E-LYLAYPAH-K] displayed on yeast cells

Recombinant biotinylated N-YAP and WW-YAP were titrated against yeast cells displaying cyclo[*E*-LYLAYPAH-*K*] or cells displaying the linear precursor (QLYLAYPAHK) to generate yeast surface display binding isotherms as previously described.^40,52^ To evaluate the binding selectivity of the yeast displayed peptides, a set of binding isotherms were also generated using biotinylated bovine serum albumin (BSA). The amount of target protein adsorbed on the surface of the yeast cells was measured via fluorescence labeling using streptavidin-phycoerythrin. The normalized fluorescent values (**Figure 4**) were treated as adsorption data and fit to a monovalent binding isotherm to estimate the binding affinity (K_D_) of the peptide ligands (**Table 1**).

**Table 1.** Values of binding affinity (K_D_) of the protein:peptide complexes obtained from yeast display titration assays via immunofluorescence labeling; N/A: not applicable; N/D: not detectable; L/B: low binding. The average K_D_ of three independent replicates plus/minus one standard error is reported.

| Peptide ligand | $K_D$ ($\mu\text{M}$) | | |
| --- | --- | --- | --- |
|  | N- YAP | WW-YAP | BSA |
| cyclo[*E*-LYLAYPAH-K] | $1.75 \pm 0.33$ | $0.68 \pm 0.21$ | N/D |
| QLYLAYPAHK | $0.74 \pm 0.15$ | L/B | N/A |

**Figure 4.**
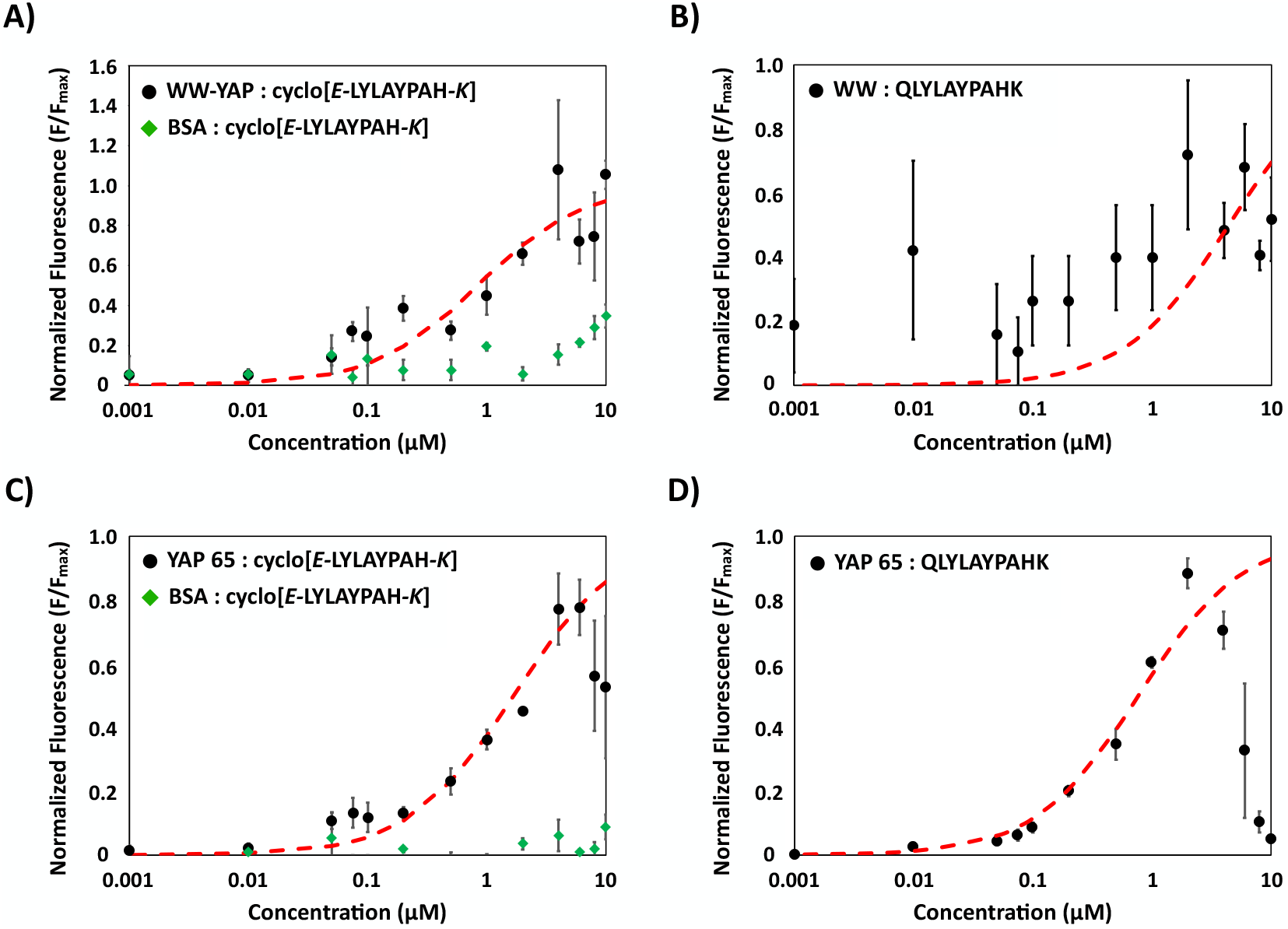
Yeast surface display binding isotherms of **(A)** WW domain of YAP (WW-YAP, black circles) and bovine serum albumin (BSA, green diamonds) on cyclo[*E*-LYLAYPAH-*K*]; **(B)** WW-YAP on linear peptide cognate QLYLAYPAHK; **(C)** N-terminal N-YAP (N-YAP, black circles) and BSA (green diamonds) on cyclo[*E*-LYLAYPAH-*K*]; and **(D)** N-YAP on QLYLAYPAHK. Yeast cell fluorescence was detected following incubation with biotinylated target protein using streptavidin-phycoerythrin conjugate. Data from each of the three independent replicates were normalized to its associated F_max_ value. The resulting values of normalized mean fluorescence intensity were fit against a monovalent binding isotherm (red dashed line). Fluorescence data from three independent experiments was used; the error bars correspond to the standard error of the mean from three independent replicates.

Most notably, the cells displaying transglutaminase-cyclized peptides exhibited a significantly higher fluorescence when titrated with WW-YAP than BSA (**Figure 4A**). Cells displaying cyclo[*E*-LYLAYPAH-*K*] returned an apparent K_D_ of 0.68 ± 0.21 μM, which is consistent with other WW domain binding peptides;^53^ the corresponding value for BSA could not be calculated, as the isotherm did not reach saturation within the tested range of protein concentration. Furthermore, the linear precursors exhibited a much lower binding affinity (**Figure 4B**), with no obvious isotherm-like binding behavior. This indicates that transglutaminase-mediated cyclization endows the peptides with specific and strong WW-binding activity.

Similarly, the apparent K_D_ of cyclo[*E*-LYLAYPAH-*K*] to YAP was 1.75 ± 0.33 μM, with a pronounced “hook” effect occurring at ~ 8 μM of YAP, as has been previously described in the case of yeast surface display titrations (**Figure 4C**).^42^ Once again, the peptide was found to bind YAP specifically, as it exhibited little-to-no BSA binding. On the other hand, the difference in binding affinity of the cyclic *vs.* linear peptide observed with WW-YAP was not observed with N-terminal YAP (**Figure 4D**). This may be imputed to additional targeting of the linear peptide to other motifs in YAP, homologous yet extraneous to the WW domain, that stabilize their interaction.

### Construction and optimization of yeast displayed cyclic peptides via intracellular transglutami-nase-mediated cyclization

Following up on the screening of yeast display libraries where peptide cyclization had been accomplished via extracellular enzymatic treatment, we also sought to explore an intracellular route to transglutaminase-mediated peptide cyclization (**Figure S1**). In the yeast display platform, the expression of peptides on the cell surface is controlled by the inducible galactose promoter Gal 1/10. The bidirectional nature of Gal 1/10 has been leveraged to achieve concurrently the display of peptides and the expression of enzymes that can edit their chemical composition and structure. For example, the tobacco etch virus protease was engineered via the Yeast Endoplasmic Sequestration System (YESS) to achieve dual display of a TEV enzyme library in conjunction with specific and non-specific selection substrates.^38^ In this approach, both enzyme and substrate transit through the endoplasmic reticulum (ER), where the enzyme can act on the substrate prior to its display on the cell surface. In another example, the modifying enzyme Proc M and its lanthipeptide substrate were co-expressed resulting in a macrocyclic lanthipeptide displayed on the yeast surface.^24^ In a similar fashion, we sought to express transglutaminase under the Gal10 promoter together with a model peptide substrate under the control of the Gal1 promoter (**Figure S2A**). After inducing the yeast cells in galactose-containing media, both transglutaminase and the peptide substrate were co-expressed as fusion proteins with their corresponding tags and signal location sequences. In addition to the standard N terminal prepro secretory signal sequence used to direct the fusion proteins toward the secretory pathway, the endoplasmic reticulum retention sequence FEHDEL^38^ was added to the C-terminus of the peptide constructs. The peptide FEHDEL, in fact, increases the residence time of the protein fusion by docking onto the ER membrane. This hybrid mechanism of intracellular trafficking and peptide cyclization is expected to ultimately afford the display of cyclic peptides only on the surface with higher cyclization yield compared to the extracellular cyclization approach.

The model lead sequence ALQSGSRGGG fused to the c-myc epitope tag EQKLISEEDL, which contains the lysine (K) residue needed for cyclization, was used to evaluate this hypothesis. The full sequence of *Streptomyces mobaraensis* transglutaminase (GenBank accession number AAT65817.1, 395 AAs) was expressed under the Gal10 promoter; the protein comprises a 60 AA-long N terminal pro sequence and a 335-AA long transglutaminase enzyme. The yield of the cyclization reaction was evaluated via immunofluorescent flow cytometry using the c-myc tag detection and the TEV protease-cleavage indirect assays described above. Upon induction at different time points (24, 48, 72, and 96 hr), the cells were treated with TEV protease to attempt the cleavage of the peptide fusion from the yeast surface. Notably, a marked difference was observed compared to the intracellular cyclization route: the treatment with TEV protease always resulted in a significant reduction in the c-myc epitope detection signal (**Figures S2B – S2E**), suggesting that most of the displayed peptides are linear.

To improve the yield of intracellular cyclization, we modified the structure of the transglutaminase fusion protein. The full-length enzyme contains a pro-peptide sequence that forms an α-helix into the catalytic site of transglutaminase and is typically cleaved by intracellular endopeptidases after translation and protein folding.^54^ We hypothesized that this α-helix remains tethered to the folded enzyme, thus hindering its active site and preventing the cyclization of the model peptide.

To this end, two different transglutaminase variants were initially explored. First, we introduced a Kex2 endopeptidase cleavage recognition site (K-R) between the pro-peptide and the transglutaminase sequence, which has been observed to promote the secretion of active transglutaminase in *Pichia pastoris* and *Candida boidinii*.^54^ Kex2 peptidase is expressed endogenously in *Saccharomyces cerevisiae* and is present in the Golgi as part of the secretory pathway.^55^ Next, we introduced a single point mutation into the substrate sequence by mutating the glutamine (Q) into an alanine (A). This mutation removes one of the two residues necessary for cyclization and enables a direct comparison between the different “intracellular” constructs. The constructs of the “Pro-Kex-TG” vector, which contains the Kex2 protease recognition sequence, and the “Pro-Kex-TG w/ Q-to-A SDM” vector, which contains both the recognition sequence and a “Q-to-A” mutation (SDM: site directed mutagenesis), are shown in **Figures S3A** and **S3B**. The cells populations were induced for 48 hrs and analyzed via immunofluorescent flow cytometry. The yeast display construct was characterized by measuring the c-myc tag level via fluorescent flow cytometry. Treatment of the “Pro-Kex-TG” construct with TEV protease caused a significant decrease in c-myc detection (**Figure 5A**), indicating cleavage from the cells, which suggests that the intracellular cyclization yield is rather modest. The corresponding characterization of the constructs where peptide cyclization was achieved via extracellular incubation with transglutaminase (**Figure 2C**) indicated that peptide cyclization reduces TEV proteolysis and the cells maintain appreciable residual levels of c-myc. These observations were corroborated by measuring the c-myc levels of “Pro-Kex-TG” and “Pro-Kex-TG w/ Q-to-A SDM” constructs. As observed above, peptide cyclization results in a decrease of the c-myc signal due to the chemical modification of the tag sequence (**Figure 2A**). The c-myc signal of the “Pro-Kex-TG” construct, however, is high and comparable to that of the “Pro-Kex-TG w/ Q-to-A SDM”, whose cyclization is prevented by the Q-to-A modification, indicating low or no cyclization (**Figure 5B**). At the same time, the HA signal is normal, which confirms that both constructs are displayed on the surface of yeast (**Figure 5B**). Representative flow cytometry plots indicate that both constructs are displayed at a detectable surface density on cells. However, there is no detectable difference in HA or c-myc tag expression in either the “Pro-Kex-TG” or “Pro-Kex-TG w/ Q-to-A SDM” construct when induced for either 24 hrs or 48 hrs (**Figure S4A**). While treatment with TEV protease does not alter the HA signal, the marked decrease in the c-myc signal in both constructs indicates low to no cyclization (**Figure S4B**). Collectively, these data indicate that the insertion of the Kex2 recognition sequence between the pro-peptide and active transglutaminase does not improve the cyclization efficiency.

**Figure 5.**
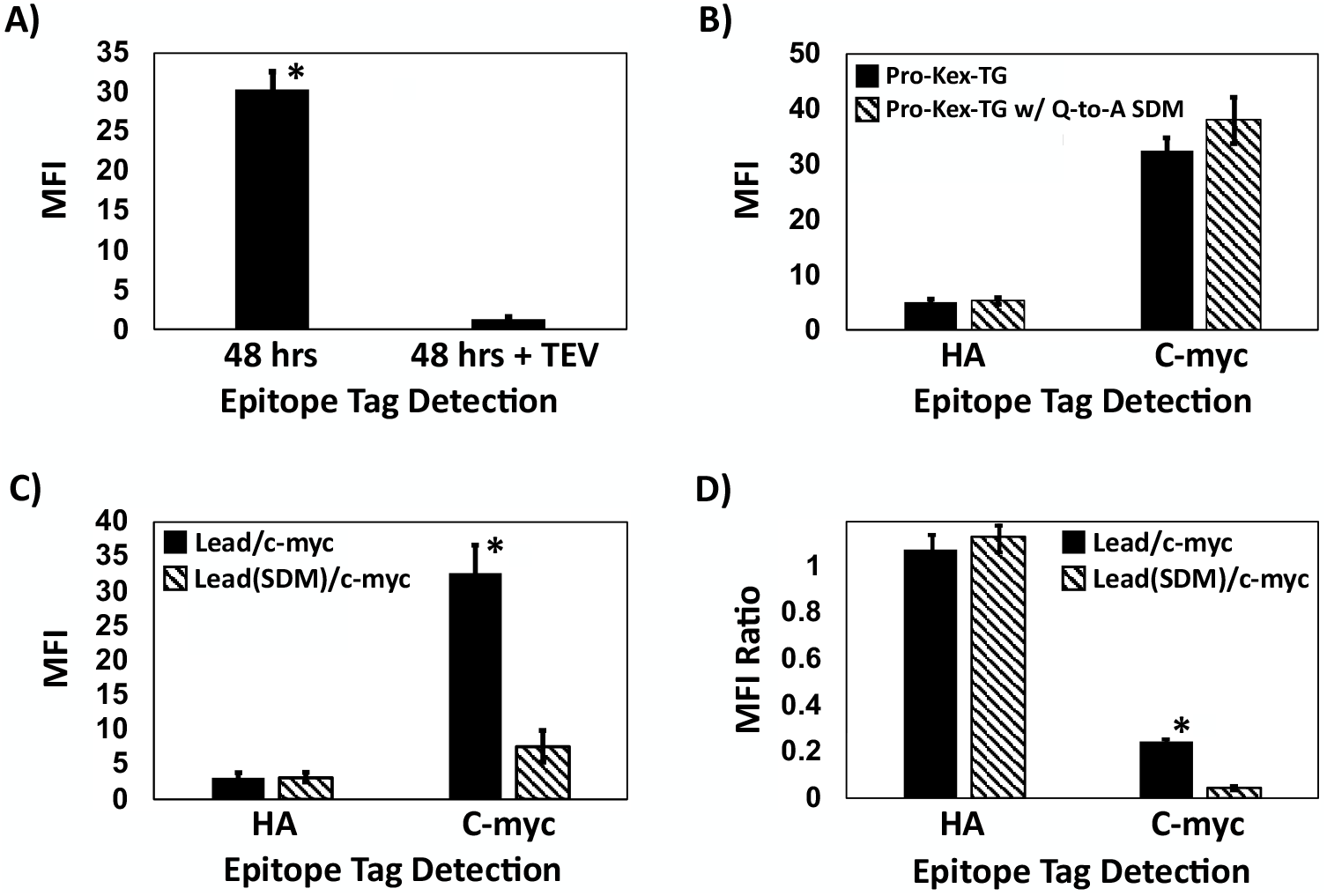
Mean fluorescence intensity (MFI) of **(A)** c-myc tag levels detected on yeast cells engineered with the “Pro-Kex-TG” vector and induced for 48 hrs before and after treatment with TEV protease; **(B)** HA and c-myc tag levels detected on yeast cells engineered with either the “Pro-Kex-TG” vector or the “Pro-Kex-TG w/ Q-to-A SDM” vector and induced for 48 hrs before and after treatment with TEV protease; and **(C)** HA and c-myc tag levels detected on yeast cells induced for 48 hrs to express either the “lead/c-myc” or the mutated “Q-to-A lead/c-myc” construct before treatment with TEV protease; **(D)** MFI ratio HA and c-myc tag levels detected on yeast cells induced for 48 hrs to express either the “lead/c-myc” or the mutated “Q-to-A lead/c-myc” construct after treatment with TEV protease (*note:* the MFI ratio is the ratio of MFI measured after *vs.* before treatment with TEV protease). The error bars correspond to the standard error of the mean from three independent replicates. A two-tailed paired t-test was performed, where * indicates p<0.05.

We therefore attempted an alternative route, which consists in expressing active transglutaminase (AAs 61-395) under the control of the Gal10 promoter concurrently with the c-myc epitope tag EQKLISEEDL in downstream to either the lead sequence ALQSGSRGGG (lead/c-myc) or the mutated sequence ALASGSRGGG (Q-to-A lead/c-myc) under the control of the Gal1 promoter (**Figures S3C** and **S3D**). Both cell populations were induced for 48 hrs, treated with TEV protease, and analyzed via immunofluorescent flow cytometry to detect the levels of HA and c-myc tags (**Figures 5C** and **5D**; representative histograms are shown in **Figures S4C** and **S4D**). As anticipated, the HA signal detected on “lead/c-myc” construct did not differ significantly upon TEV protease treatment (**Figure 5D**). The comparison between the c-myc tag levels of the construct “lead/c-myc” *vs.* “Q-to-A lead/c-myc” (**Figure 5C**) and “Pro-Kex-TG” (**Figure 5A**) prior to treatment with TEV protease suggests that peptide cyclization has occurred on the “lead/c-myc” construct to some extent. The fluorescent signal obtained with the “lead/c-myc” construct, in fact, is 4-fold lower than that of the other two constructs and is comparable with that observed with peptide cyclized via extracellular treatment with transglutaminase (**Figure 2A**). Nonetheless, the decrease in c-myc tag signal with the “lead/c-myc” construct upon TEV treatment indicates that a significant fraction of the yeast displayed peptides had not been cyclized (**Figure 5C**); comparatively, however, the loss of c-myc signal for the “Q-to-A lead/c-myc” construct upon treatment with TEV protease was much stronger (**Figure 5D**); the Q-to-A mutation completely prevents peptide cyclization, allowing TEV proteolysis to proceed unhindered. Observing a bigger loss of c-myc level upon TEV proteolysis with the linear construct compared to the “Active TG” construct corroborates the claim that the latter has undergone partial cyclization.

Notably, conclusive differences in fluorescence signal were observed only with cells induced for 48 hrs, whereas little-to-no variation was observed with cells induced for 48 hrs (**Figures S4D**). This suggests the need for a sufficiently high level of transglutaminase in the ER to achieve detectable peptide cyclization. The addition of a ER-docking sequence in downstream to the transglutaminase sequence promotes its interaction with the peptide substrate. The enzyme is ultimately secreted into the supernatant, where it may catalyze additional (extracellular) cyclization of the peptides displayed on the Aga2 systems. Collectively, these results show that the co-expression of active TG and the substrate peptide construct enable the cyclization of the peptide on the surface of the yeast cells.

### Construction and screening of yeast display libraries of cyclic peptides via intracellular transglutaminase-mediated cyclization and characterization of the identified peptides

Upon optimization of the transglutaminase sequence and the induction time, we sought to utilize this design to construct yeast display libraries of cyclic peptides to be screened for the identification of YAP-binding ligands. Accordingly, the linear heptapeptide library ALQX_1_X_2_X_3_X_4_X_5_X_6_X_7_EQKLISEEDL was constructed, wherein the length and design of both leader and c-myc tag segments were maintained as in the optimized construct to ensure high cyclization efficiency. The library was screened against N-terminal YAP by conducting one round of magnetic cell sorting and one of FACS. While little enrichment was observed after magnetic selection, a marked selection towards YAP binding was observed after FACS (**Figure S5**). Of the five clones selected for sequencing, two returned the sequence cyclo[*E*-VQCRGKGEQ-*K*], whereas the other three comprised sequences that were not consistent with the original construct (*i.e.*, lacking the leader sequence or out of frame) and were discarded. The YAP:cyclo[*E*-VQCRGKGEQ-*K*] interaction, evaluated via yeast surface titration assays, returned an apparent K_D_ of 0.46 ± 0.08 μM. To evaluate the peptide’s binding specificity to YAP, yeast surface titrations were executed against biotinylated BSA. The comparison of the YAP *vs.* BSA isotherms (**Figure 6A**) indicates that the selected peptide is a selective ligand. To assess the binding interaction of linear AVQCRGKGEQK with YAP, a similar mutation strategy was carried out to convert the reactive glutamine (Q) residue into an unreactive alanine (A) thus leading to the surface display peptide sequence AVQCRGKGEQK. The YAP:AVQCRGKGEQK interaction, evaluated via yeast surface titration assays, returned an apparent K_D_ of 0.36 ± 0.07 μM (**Figure 6B**). The apparent similarity in YAP-binding strength between the linear and cyclic peptide could be imputed to incomplete cyclization of the peptides in the yeast display library. The screening of yeast cells displaying a combination of cyclic and linear peptides may in fact lead to the identification of peptide sequences that bind the target protein in both configurations and with comparable affinity. While optimization is needed, these proof-of-concept data show the potential of yeast display libraries of peptides cyclized intracellularly via transglutaminase-mediated amidation as effective tools for the rapid discovery of ligands with selective biorecognition activity towards protein targets. Since TEV treatment is likely to eliminate non-cyclized peptides from the yeast surface (**Figure 2**), one strategy for selective enrichment of cyclic peptide binders may involve TEV treatment of the library prior to combinatorial screening.

**Figure 6.**
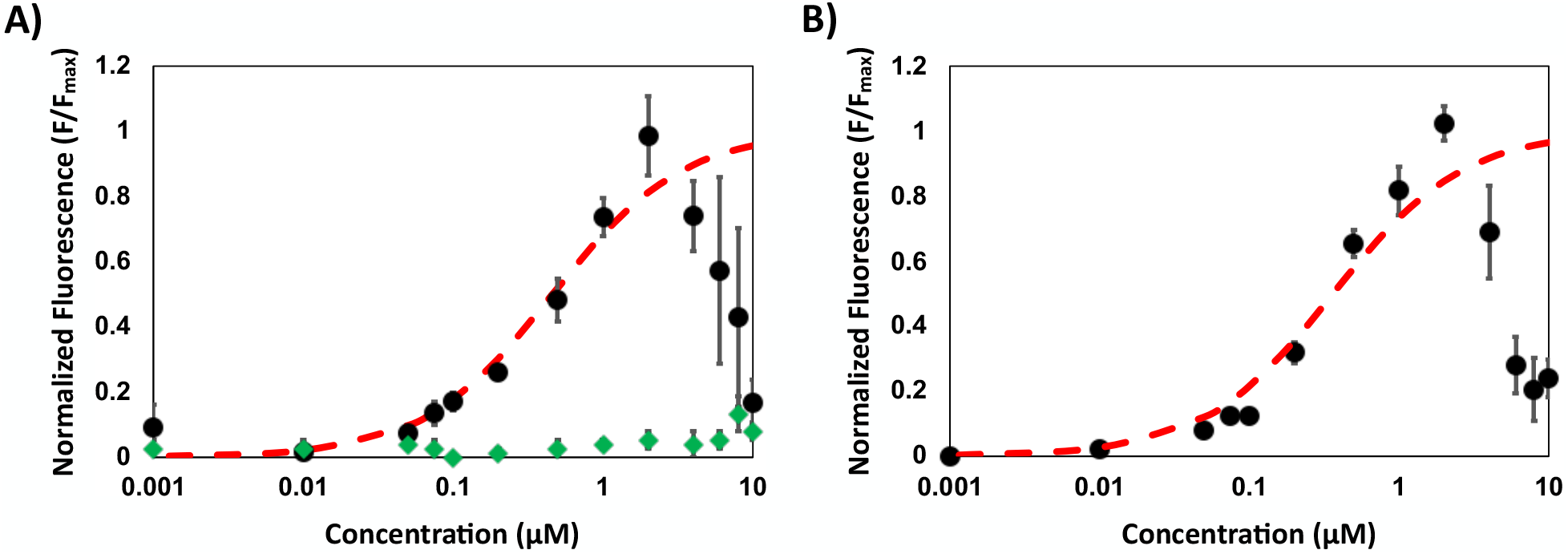
Yeast surface display binding isotherms of N-terminal N-YAP (N-YAP, black circles) and BSA (green diamonds) on **(A)** cyclo[*E*-VQCRGKGEQ-*K*] and **(B)** its linear counterpart QVQCRGKGEQK. Yeast cell fluorescence was detected following incubation with biotinylated target protein using streptavidin-phycoerythrin conjugate. Data from each independent replicate was normalized to its associated F_max_ value. The resulting values of average normalized mean fluorescence intensity were fit against a monovalent binding isotherm (red dashed line). Fluorescence data from three independent experiments was used; the error bars correspond to the standard error of the mean from three independent replicates.

### Binding affinity and specificity of cyclo[E-LYLAYPAH-K] conjugated to chromatographic resin

Our results using yeast surface titrations showed that cyclo[*E*-LYLAYPAH-*K*] showed selective binding to WW-YAP and bound with higher affinity than its linear counterpart. We further evaluated the biorecognition activity of cyclo[*E*-LYLAYPAH-*K*] as a synthetic construct on solid phase. To this end, the peptide sequence, where an N-terminal glutamic acid (E) was used instead of glutamine (Q), was synthesized on Toyopearl amino resin and cyclized by reacting the γ-carboxylic acid of E with the ε-amino group of K. This tethering strategy makes the cyclic synthesized peptide chemically identical to that obtained by transglutaminase-mediated cyclization. The known peptide sequence RYSPPPPYSSHS comprising of the PPxY-like WW-binding domain within the PTCH peptide was also conjugated on Toyopearl resin to serve as positive control. Protein adsorption isotherms were generated by incubating aliquots of peptide-Toyopearl resin in solutions of either N-terminal YAP or WW-YAP at different concentrations ranging between 20 μg/mL and 3 mg/mL. The adsorption data (*i.e.*, the amounts of protein adsorbed on solid phase *vs.* the corresponding values of protein concentration in solution at the binding equilibrium) of WW-YAP to cyclo[*E*-LYLAYPAH-*K*]-Toyopearl and PTCH-Toyopearl resins are reported in **Figure 7A** and **7B**, respectively; the corresponding binding isotherms of N-terminal YAP are in **Figure 7C** and **7D**, respectively.

**Figure 7.**
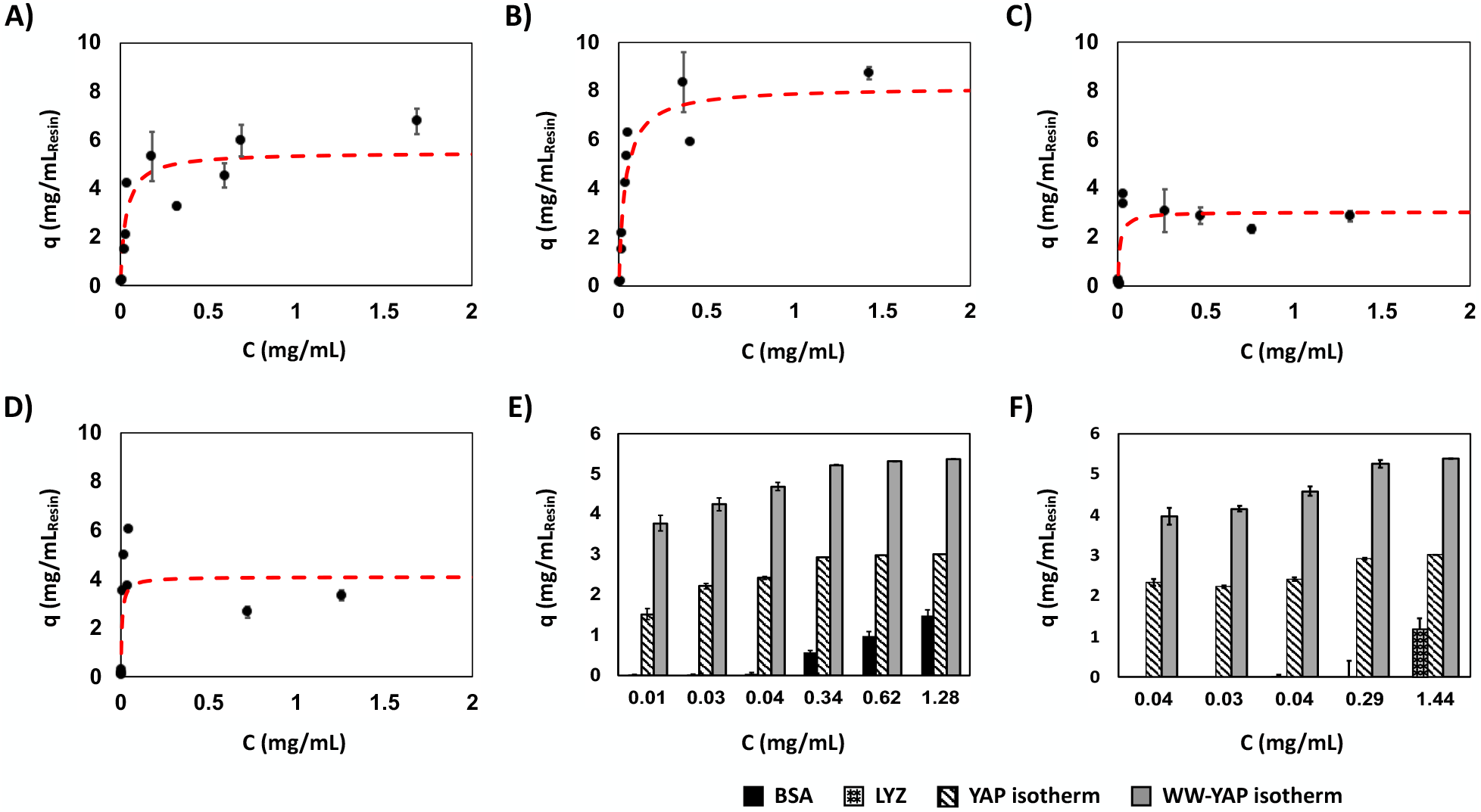
Binding isotherms of (A) WW-YAP on RYSPPPPYSSHS-Toyopearl resin; **(B)** WW-YAP on cyclo[*E*-LYLAYPAH-*K*]-Toyopearl resin; (A) N-YAP on RYSPPPPYSSHS-Toyopearl resin; **(B)** N-YAP on cyclo[*E*-LYLAY-PAH-*K*]-Toyopearl resin; **(E)** Comparison of experimental data of bovine serum albumin (BSA) adsorption compared to corresponding values of WW and YAP adsorption derived from the isotherms **(B)** and **(D)** at the same values of equilibrium protein concentration; **(F)** Comparison of experimental data of lysozyme (LYZ) adsorption compared to corresponding values of WW and YAP adsorption derived from the isotherms **(B)** and **(D)** at the same values of equilibrium protein concentration. The values of protein adsorption were determined from three independent experiments; the error bars correspond to the standard error of the mean from three independent replicates. A two-tailed t-test assuming unequal variances was performed at each concentration to compare the adsorbed amount of lysozyme or BSA to the isotherm-calculated values; the differences were statistically significant with p<0.05.

The values of K_D_ and Q_max_ obtained by fitting the adsorption data to a Langmuir binding isotherm are collated in **Table 2**. Notably, the values of apparent K_D_ of cyclo[*E*-LYLAYPAH-*K*] are an order of magnitude lower (*i.e.*, higher binding strength) than those obtained via yeast surface display titration assays. This can be imputed to the much higher value of surface density of peptide ligands conjugated on Toyopearl resin (~ 7 μmol/m^2^) compared to corresponding value on the surface of yeast cells (~ 2 nmol/m^2^). It is also noticed that while the adsorption isotherms of WW-YAP on both the cyclic peptide and the control PTCH peptide feature a standard monovalent binding isotherm shape, those obtained with N-terminal YAP are rather irregular. Specifically, both isotherms display the characteristic rapid increase in YAP binding within the range of equilibrium protein concentration below 0.5 mg/mL; however, as the latter approaches and exceeds ~ 1 mg/mL, the isotherms do not feature the characteristic plateau, but exhibit a slight decrease in the amount of adsorbed protein. This suggests that the layer of proteins adsorbed on the peptide-functionalized surface of the chromatographic resin may be unstable, and the free YAP in solution may interfere with the YAP:peptide complexes and trigger their dissociation.

**Table 2.** Values of binding affinity (KD) and capacity (Q_max_) of peptide-Toyopearl resins for the WW domain of YAP (WW-YAP) and N-terminal YAP obtained by fitting the adsorption data in **Figure 7** with Langmuir isotherms.The average K_D_ of three independent replicates plus/minus one standard error is reported.

| Peptide ligand | Target Protein |  |  |  |
| --- | --- | --- | --- | --- |
|  | WW-Yap |  | N-terminal YAP |  |
| | $K_D$ (μM) | $Q_{max}$ (mg/mL) | $K_D$ (μM) | $Q_{max}$ (mg/mL) |
| cyclo[E-LYLAYPAH-K] | $2.64 \pm 0.71$ | $5.60 \pm 0.42$ | $0.34 \pm 0.02$ | $3.03 \pm 0.13$ |
| PTCH peptide | $2.88 \pm 0.48$ | $8.3 \pm 0.27$ | $0.20 \pm 0.07$ | $4.11 \pm 0.12$ |

We also evaluated the binding specificity of resin-bound cyclo[*E*-LYLAYPAH-*K*] using bovine serum albumin (BSA) and lysozyme (LYZ) as negatively and positively charged control proteins, respectively. The peptide-Toyopearl resin were incubated with solutions of BSA or LYZ in PBS at concentrations ranging from 13 μg/mL to 2 mg/mL. The resulting adsorption data were compared to the corresponding amounts of bound WW-YAP or N-terminal YAP derived from the isotherms at the same values of equilibrium protein concentration in solution. These comparisons, summarized in **Figure 7E** for BSA and **Figure 7F** for LYZ, show significantly lower binding of both control proteins, demonstrating the specific biorecognition activity of cyclo[*E*-LYLAYPAH-*K*] for the targets WW-YAP and N-terminal YAP.

Collectively, these results show that cyclic peptide binders isolated from transglutaminase-cyclized yeast display libraries exhibit target binding and selectivity when chemically synthesized on chromatographic resins.

## Conclusions

In this work, we have demonstrated the use of enzymatic modification by transglutaminase as an efficient route to produce yeast display libraries of cyclic peptides. To this end, we have developed two routes, one based on extracellular and one on intracellular transglutaminase-mediated amidation, and for each tailored the design of the yeast display Aga2-based constructs to optimize the yield of peptide cyclization. The integration of epitope detection tags, such as the HA and the c-myc tags, and cleavage sites, such as the TEV protease recognition sequence, allowed us to assess the degree of cyclization at the level of both single-sequence and library. While the peptide cyclization via intracellular catalytic amidation afforded a relatively low yield, its extracellular counterpart performed with soluble transglutaminase proved significantly more efficient. Nonetheless, important findings were made on how the design of the Aga2-based peptide construct impact – and improve – the cyclization yield. In this regard, future work could focus on improving the design presented herein and accomplish the cyclization of most or all of the peptides displayed on the surface of yeast cells. Regarding screening, the implementation of magnetic selection and FACS enabled the rapid identification of cyclic peptide affinity ligands with selective biorecognition for WW-YAP and N-terminal YAP. The ligand discovery and characterization methods developed in this work are agnostic to the specific peptide cyclization chemistry and can be expanded to other enzyme-mediated reactions. Collectively, this makes the work presented here of wide interest towards the rapid development of biorecognition agents for modulating biochemical processes, as well as detection and purification of target proteins. In particular, the apparent binding affinities of cyclic peptides isolated using our approach (micromolar as estimated by yeast titrations) make them well suited for use in chromatography applications.

## Acknowledgements

This work was funded by the National Science Foundation (CBET 1510845). JB kindly acknowledge support from the National Institute of Health Molecular Biotechnology Training Fellowship (NIH T32 GM008776). The authors also wish to thank the UNC High-Throughput Peptide Synthesis and Array facility for peptide synthesis and the UNC Flow Cytometry Core facility for use of their FACSAria II (Becton Dickinson).

## Electronic Supplementary Information

**Table S1.**
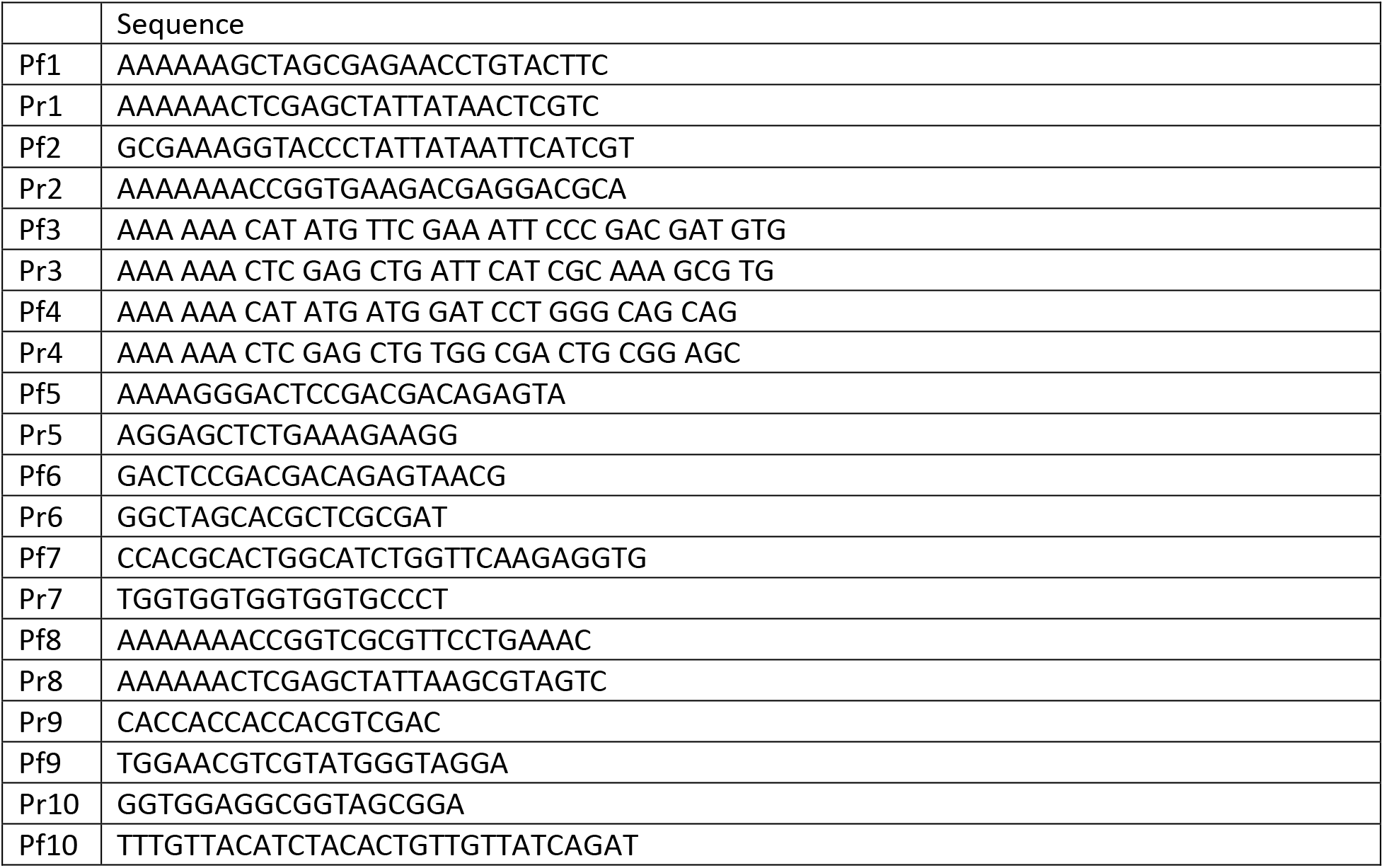
Oligonucleotide primers.

**Table S2.**
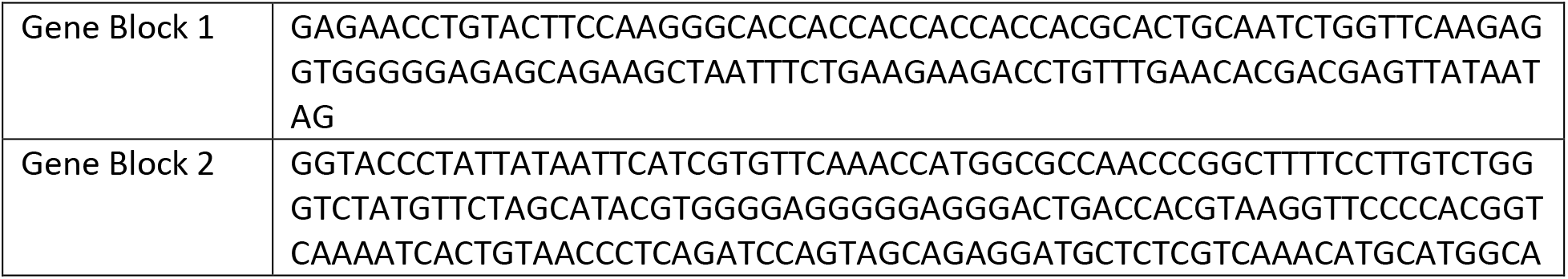
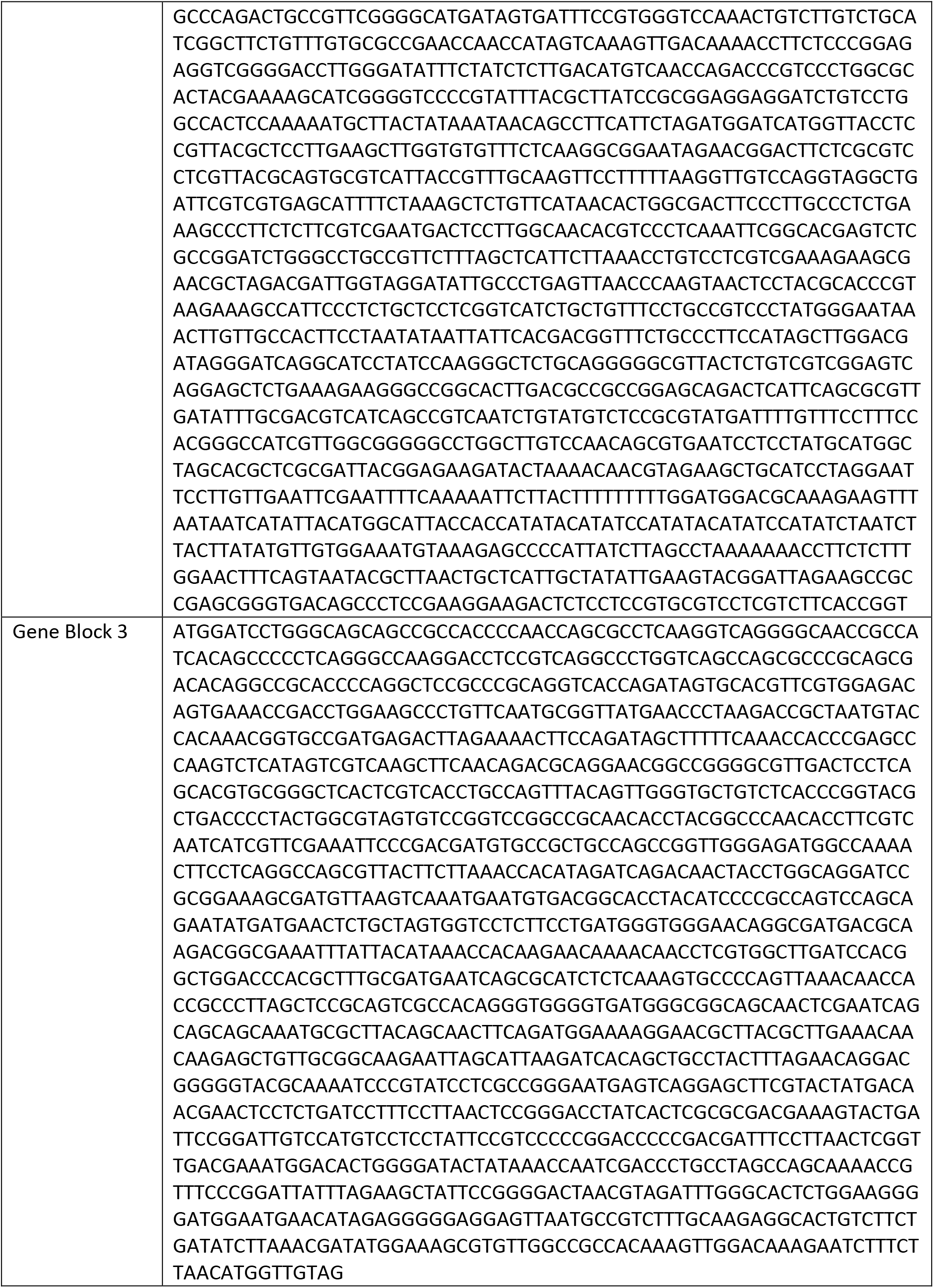
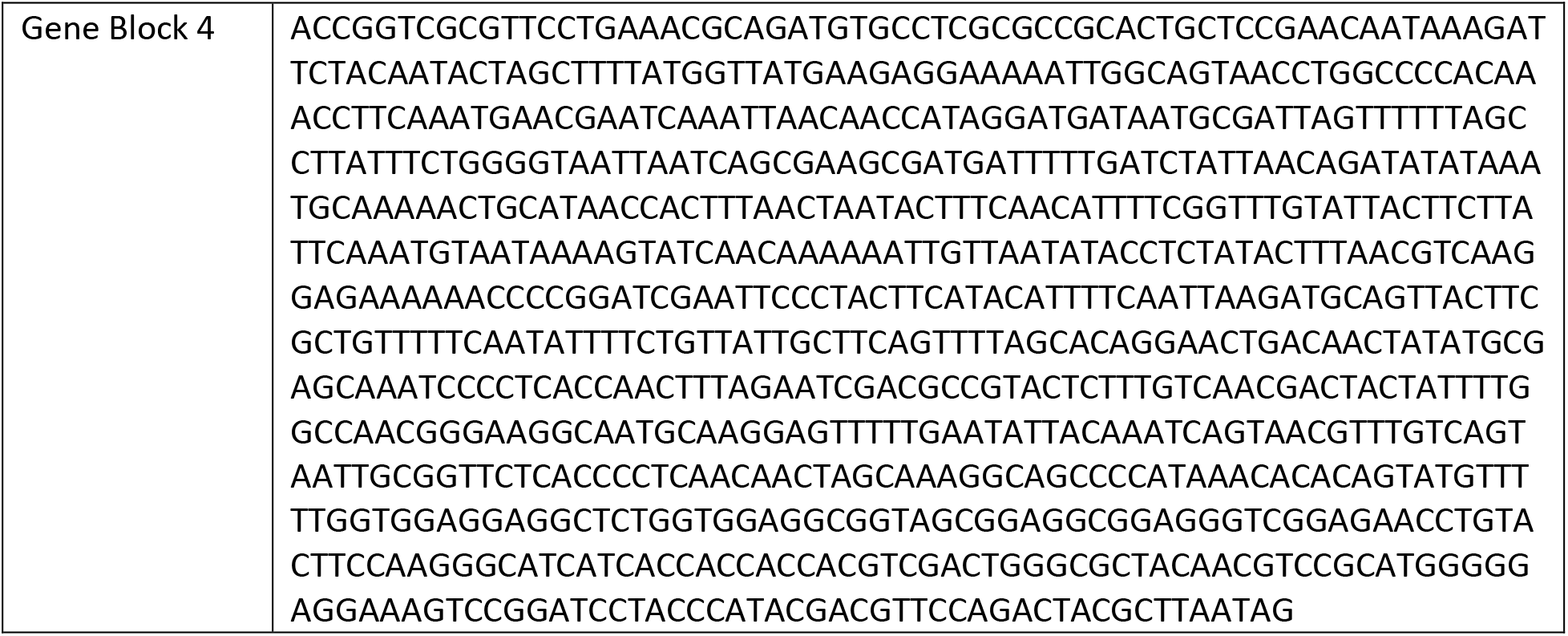
Gene Blocks.

**Table S3.**
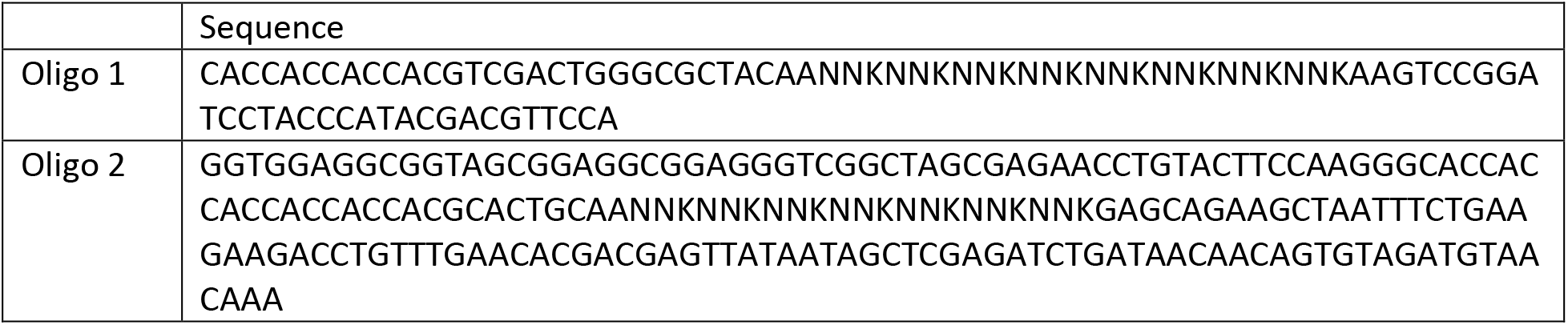
Oligonucleotides encoding randomized peptides.

**Figure S1.**
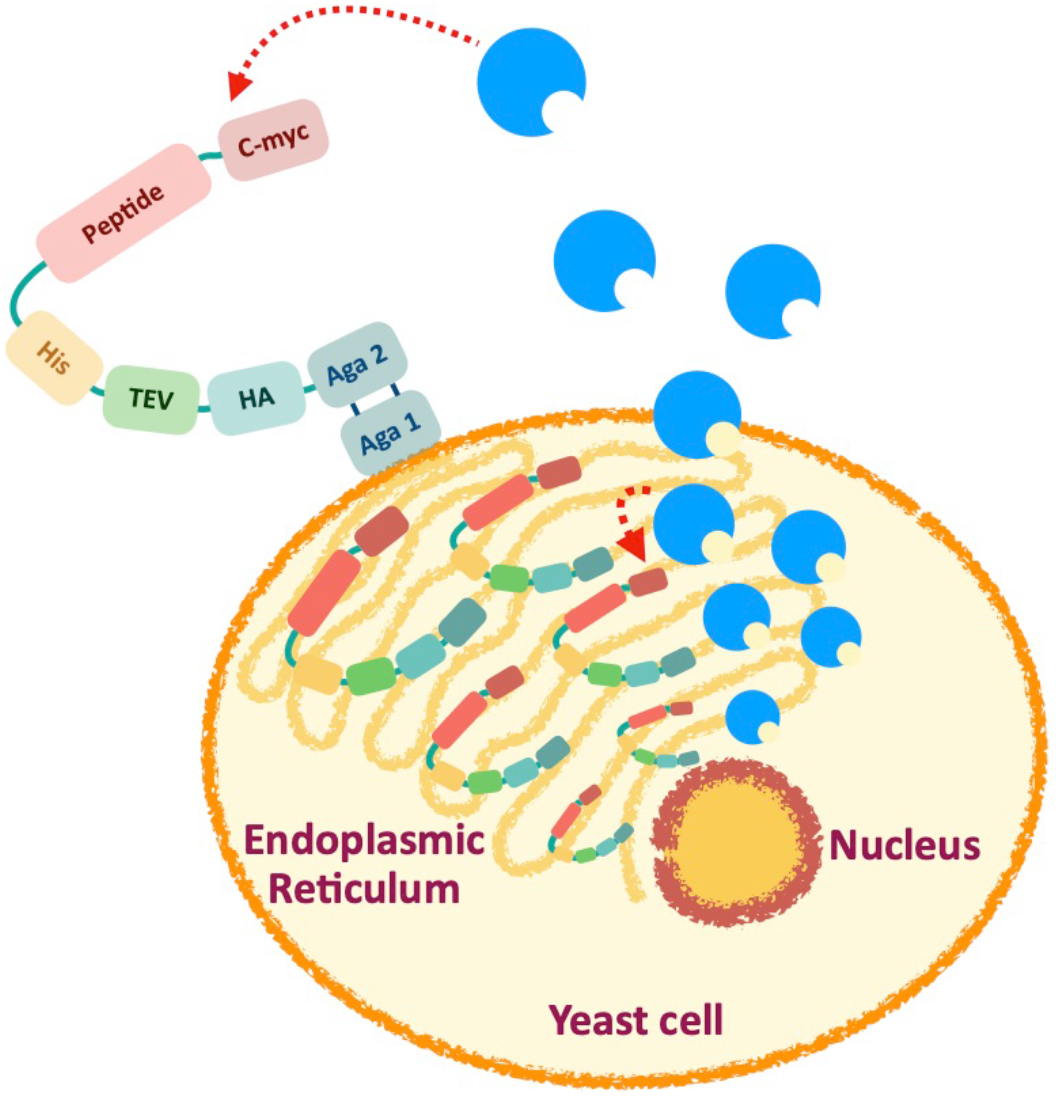
Co-expression of the peptide construct and transglutaminase to achieve the cyclization of the model peptide either intracellularly, during trafficking through the endoplasmic reticulum, or upon display on the surface of the yeast cell.

**Figure S2.**
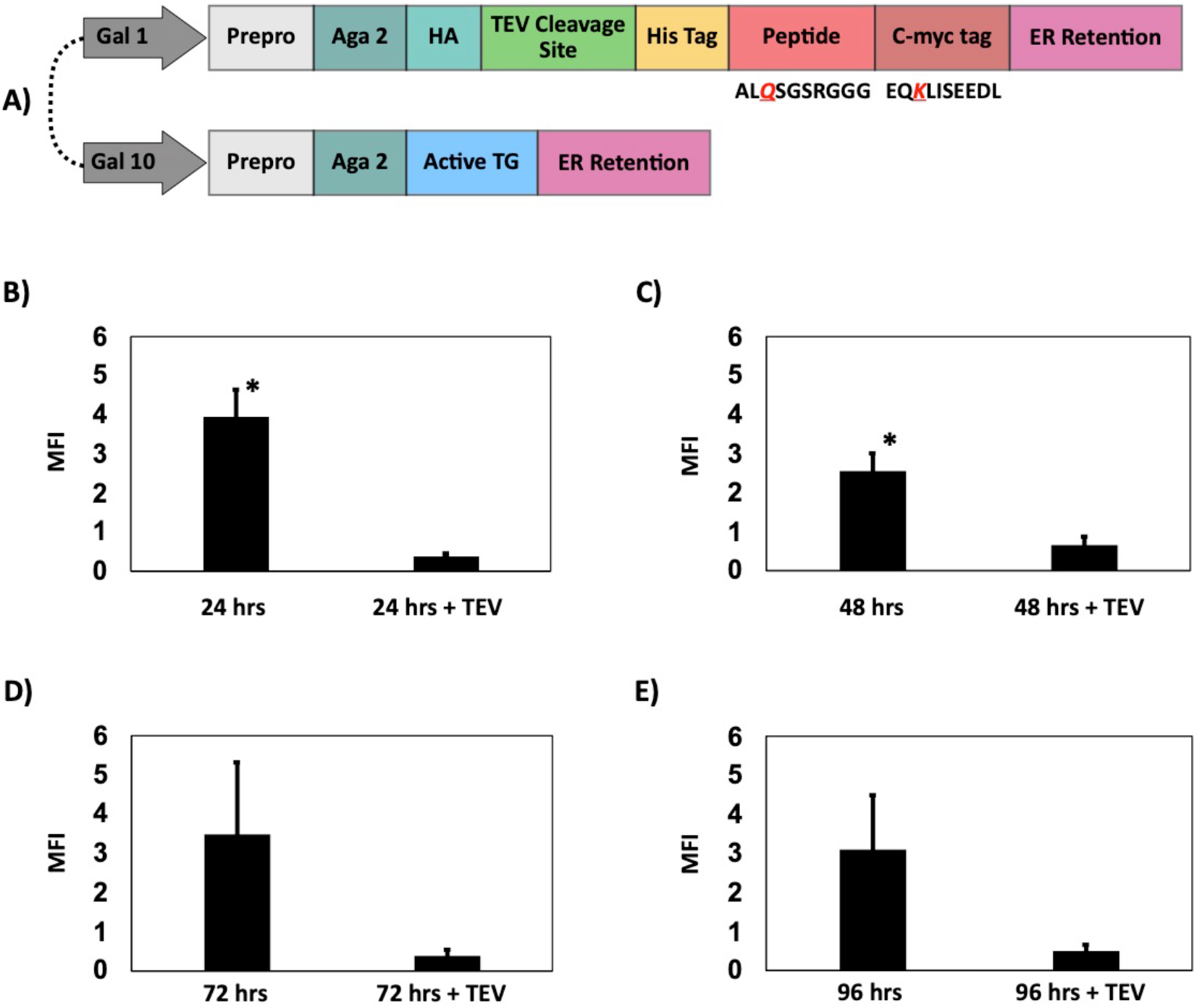
**(A)** Design of the pCTcon2 plasmid vector with bidirectional Gal1/10 promoter to express the model peptide sequence ALQSGSRGGG fused to the c-myc epitope sequence EQKLISEEDL concurrently with the full length transglutaminase enzyme. Mean fluorescence intensity (MFI) of c-myc tag level detected before and after treatment of yeast cells with TEV protease, wherein the peptide construct and the full length transglutaminase enzyme were co-expressed for **(A)** 24 hrs, **(B)** 48 hrs, **(C)** 72 hrs, and **(D)** 96 hrs. The error bars correspond to the standard error of the mean from three independent replicates. A two-tailed paired t-test was performed; * indicates p<0.1 and ** indicates p<0.05.

**Figure S3.**
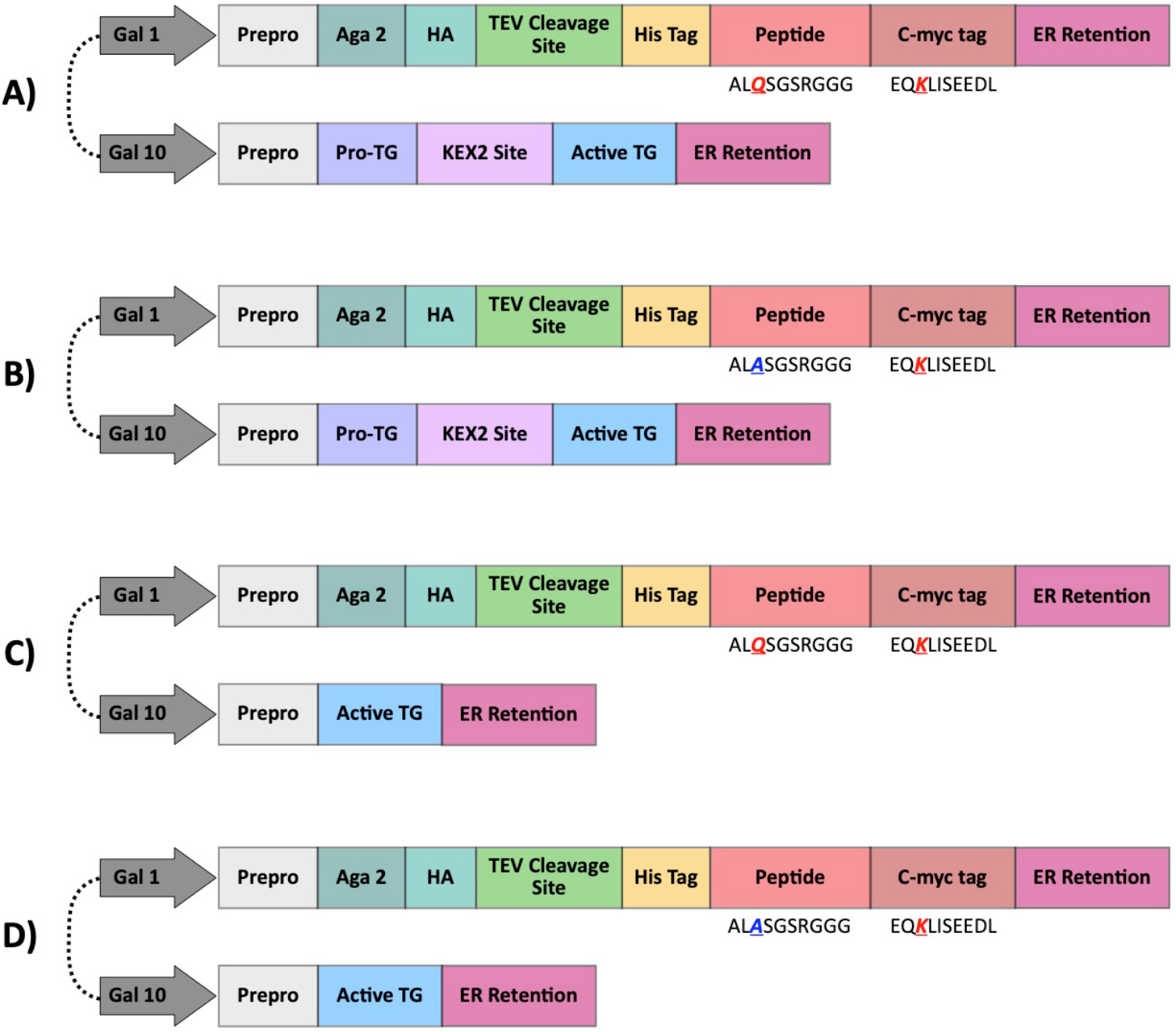
**(A)** Design of pCTcon2 plasmid vectors with bidirectional Gal1/10 promoter to express **(A)** the model peptide sequence ALQSGSRGGG fused to the c-myc epitope sequence EQKLISEEDL concurrently with a fusion protein comprising transglutaminase framed between the Kex2 endopeptidase recognition site (K-R) and the ER retention peptide; **(B)** the mutated model peptide sequence ALASGSRGGG fused to the c-myc epitope concurrently with a fusion protein comprising transglutaminase framed between the Kex2 endopeptidase recognition site and the ER retention peptide; **(C)** the model peptide sequence fused to the c-myc epitope sequence concurrently with active transglutaminase; and **(D)** the mutated peptide sequence fused to the c-myc epitope sequence concurrently with active transglutaminase.

**Figure S4.**
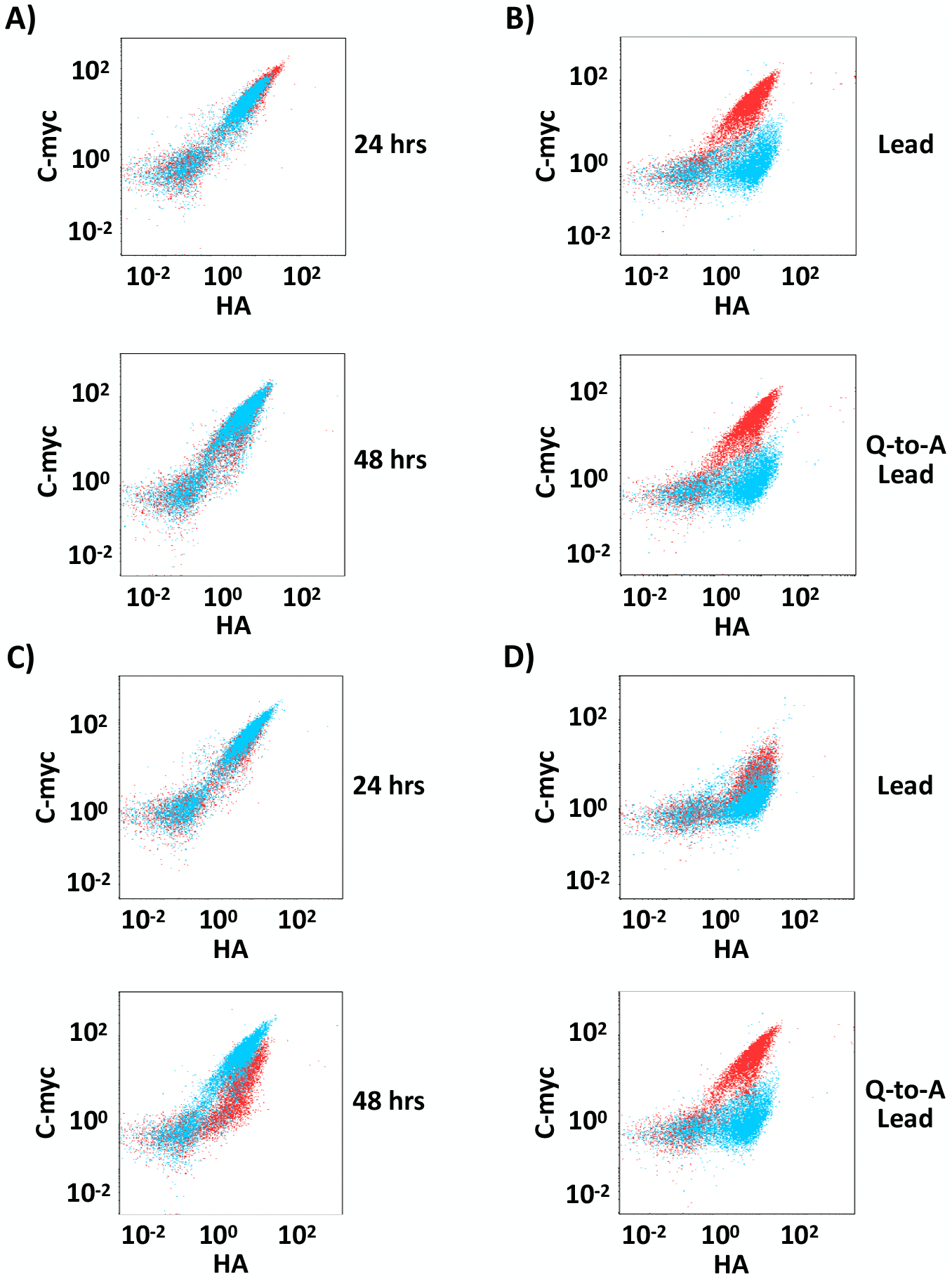
Representative flow cytometry plots comparing the c-myc tag (y-axis) and HA tag (x-axis) detection signal of **(A)** yeast cells engineered with either the “Pro-Kex-TG” vector (red) or the “Pro-Kex-TG w/ Q-to-A SDM” vector (cyan) and induced for either 24 or 48 hrs; **(B)** yeast cells engineered with either the “Pro-Kex-TG” vector or the “Pro-Kex-TG w/ Q-to-A SDM” vector and induced for 48 hrs before (red) or after (cyan) treatment with TEV protease; **(C)** yeast cells engineered to express either the “lead/c-myc” construct (red) or the “Q-to-A lead/c-myc” construct (blue) concurrently with active transglutaminase and induced for either 24 or 48 hrs; and **(D)** yeast cells engineered to express either the “lead/c-myc” construct (red) or the “Q-to-A lead/c-myc” construct (blue) concurrently with active transglutaminase and induced for 48 hrs before (red) or after (cyan) treatment with TEV protease.

**Figure S5.**
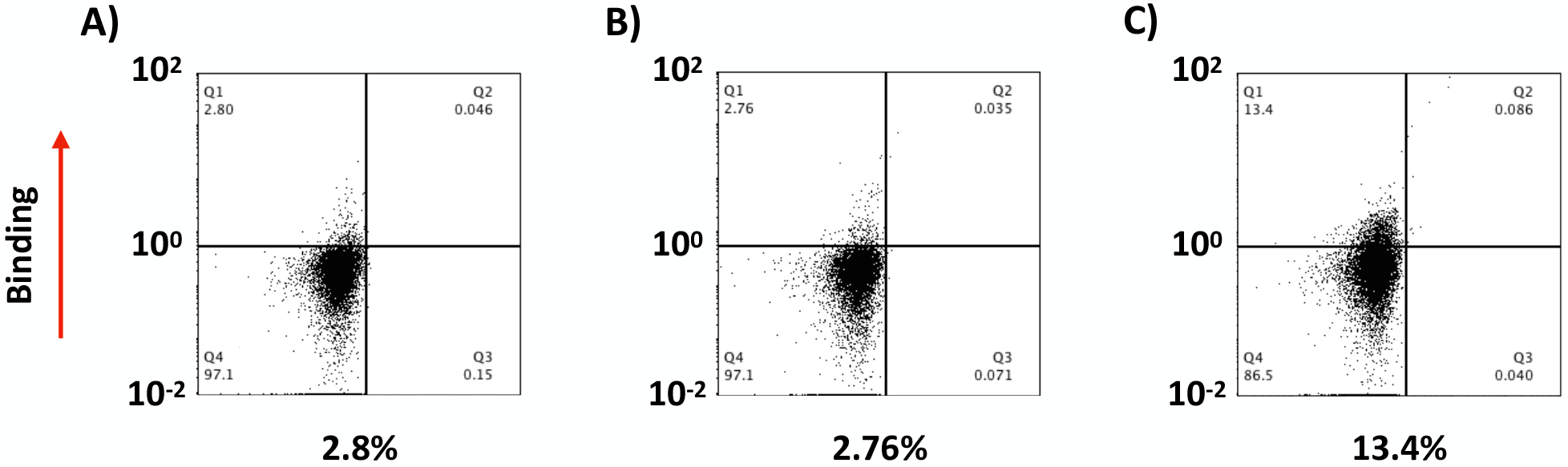
Screening of a yeast display library of peptides cyclized by intracellular transglutaminase. Flow cytometry plot of **(A)** the naïve library, **(B)** the library after one round of magnetic cell sorting, and **(C)** library after one round of magnetic cell sorting and one round of FACS. Cells were labeled with 500 nM of biotinylated YAP protein and streptavidin-phycoerythrin conjugate. The values of enrichment are reported below each plot.

## References

1. Hacker, D. E., Hoinka, J., Iqbal, E. S., Przytycka, T. M. & Hartman, M. C. T. Highly Constrained Bicyclic Scaffolds for the Discovery of Protease-Stable Peptides via mRNA Display. ACS Chem. Biol. 12, 795–804 (2017).

2. Howell, S. M. et al. Serum stable natural peptides designed by mRNA display. Sci. Rep. 4, 6008 (2014).

3. Hacker, D. E., Hoinka, J., Iqbal, E. S., Przytycka, T. M. & Hartman, M. C. T. Highly Constrained Bicyclic Scaffolds for the Discovery of Protease-Stable Peptides via mRNA Display. ACS Chem. Biol. 12, 795–804 (2017).

4. Fiacco, S. V et al. Directed Evolution of Scanning Unnatural-Protease-Resistant (SUPR) Peptides for in Vivo Applications. ChemBioChem 17, 1643–1651 (2016).

5. Middendorp, S. J. et al. Peptide Macrocycle Inhibitor of Coagulation Factor XII with Subnanomolar Affinity and High Target Selectivity. J. Med. Chem. 60, 1151–1158 (2017).

6. Millward, S. W., Fiacco, S., Austin, R. J. & Roberts, R. W. Design of Cyclic Peptides That Bind Protein Surfaces with Antibody-Like Affinity. ACS Chem. Biol. 2, 625–634 (2007).

7. Pei, D. Efficient Delivery of Cyclic Peptides into Mammalian Cells with Short Sequence Motifs. ACS Chem. Biol. 8, 423–431 (2012).

8. Bowen, J., Schloop, A., Reeves, G., Menegatti, S. & Rao, B. Discovery of membrane-permeating cyclic peptides via mRNA display. (2020). doi:10.1101/2020.07.20.212142

9. Arkin, M. R., Tang, Y. & Wells, J. A. Small-molecule inhibitors of protein-protein interactions: Progressing Towards the Reality. Chem. Biol. 18, 1102–1114 (2014).

10. Wójcik, P. & Berlicki, Ł. Peptide-based inhibitors of protein-protein interactions. Bioorg. Med. Chem. Lett. 26, 707–713 (2016).

11. Roberts, R. W. & Szostak, J. RNA-peptide fusions for the in vitro selection of peptides. Proc. Natl. Acad. Sci. 94, 12297–12302 (1997).

12. Smith, G. P. Filamentous Fusion Phage: Novel Expression Vectors that Display Cloned Antigens on the Virion Surface. Science (80-.). 228, 1315–1317 (1985).

13. Boder, E. T. & Wittrup, K. D. Yeast surface display for screening combinatorial polypeptide libraries. Nat. Biotechnol. 15, 553–557 (1997).

14. Lam, K. I. T. S., Liu, R., Miyamoto, S., Lehman, A. L. & Tuscano, J. M. Applications of One-Bead One-Compound Libraries and Chemical Microarrays in Signal Transduction Research. Acc. Chem. Res. 36, 370–377 (2003).

15. Smith, J. M., Frost, J. R. & Fasan, R. Emerging Strategies to Access Peptide Macrocycles from Genetically Encoded Polypeptides. J. Org. Chem. 78, 3525–3531 (2013).

16. Simonetti, L. & Ivarsson, Y. Genetically Encoded Cyclic Peptide Phage Display Libraries. ACS Cent. Sci. 6, 336–338 (2020).

17. Kawakami, T. & Murakami, H. Genetically Encoded Libraries of Nonstandard Peptides. J. Nucleic Acids 713510, (2012).

18. Heinis, C. Bicyclic Peptide Antagonists Derived from Genetically Encoded Combinatorial Libraries. Chimia (Aarau). 65, 677–679 (2011).

19. Zorzi, A., Deyle, K. & Heinis, C. Cyclic peptide therapeutics: past, present and future. Curr. Opin. Chem. Biol. 38, 24–29 (2017).

20. Schlippe, Y. V. G., Hartman, M. C. T., Josephson, K. & Szostak, J. W. In Vitro Selection of Highly Modified Cyclic Peptides That Act as Tight Binding Inhibitors. J. Am. Chem. Soc. 134, 10469–10477 (2012).

21. Menegatti, S., Hussain, M., Naik, A. D., Carbonell, R. G. & Rao, B. M. mRNA Display Selection and Solid-Phase Synthesis of Fc-Binding Cyclic Peptide Affinity Ligands. Biotechnol. Bioeng. 110, 857–870 (2013).

22. Bertoldo, D. et al. Phage Selection of Peptide Macrocycles against β-Catenin to Interfere with Wnt Signaling. ChemMedChem 11, 834–839 (2016).

23. Zorzi, A., Middendorp, S. J., Wilbs, J., Deyle, K. & Heinis, C. Acylated heptapeptide binds albumin with high affinity and application as tag furnishes long-acting peptides. Nat. Commun. 8, 16092 (2017).

24. Hetrick, K. J., Walker, M. C. & Van Der Donk, W. A. Development and Application of Yeast and Phage Display of Diverse Lanthipeptides. ACS Cent. Sci. 4, 458–467 (2018).

25. Palei, S., Becher, K. S., Nienberg, C., Jose, J. & Mootz, H. D. Bacterial Cell-Surface Display of Semisynthetic Cyclic Peptides. ChemBioChem 20, 72–77 (2019).

26. Tavassoli, A. SICLOPPS cyclic peptide libraries in drug discovery. Curr. Opin. Chem. Biol. 38, 30–35 (2017).

27. Osher, E. L. & Tavassoli, A. Intracellular Production of Cyclic Peptide Libraries with SICLOPPS. Methods Mol. Biol. 1495, 27–39 (2017).

28. Miranda, E. et al. A Cyclic Peptide Inhibitor of HIF −1 Heterodimerization That Inhibits Hypoxia Signaling in Cancer Cells. J. Am. Chem. Soc. 135, 10418–10425 (2013).

29. Tavassoli, A. & Benkovic, S. J. Split-intein mediated circular ligation used in the synthesis of cyclic peptide libraries in E. coli. Nat. Protoc. 2, 1126–1133 (2007).

30. Nuijens, T., Toplak, A., Schmidt, M., Ricci, A. & Cabri, W. Natural Occurring and Engineered Enzymes for Peptide Ligation and Cyclization. Front. Chem. 7, 829 (2019).

31. Schmidt, M., Toplak, A., Quaedflieg, P. J. L. M. & Nuijens, T. Enzyme-mediated ligation technologies for peptides and proteins. Curr. Opin. Chem. Biol. 38, 1–7 (2017).

32. Schmidt, M. et al. Efficient Enzymatic Cyclization of Disulfide-Rich Peptides by Using Peptide Ligases. ChemBioChem 20, 1524–1529 (2019).

33. Xu, S., Zhao, Z. & Zhao, J. Recent advances in enzyme-mediated peptide ligation. Chinese Chem. Lett. 29, 1009–1016 (2018).

34. Antos, J. M. et al. A Straight Path to Circular Proteins. J. Biol. Chem. 284, 16028–16036 (2009).

35. Nguyen, G. K. T. et al. Butelase 1: A Versatile Ligase for Peptide and Protein Macrocyclization. J. Am. Chem. Soc. 137, 15398–15401 (2015).

36. Flühe, L. et al. The radical SAM enzyme AlbA catalyzes thioether bond formation in subtilosin A. Nat. Chem. Biol. 8, 350–357 (2012).

37. Touati, J., Angelini, A., Hinner, M. J. & Heinis, C. Enzymatic cyclisation of peptides with a transglutaminase. ChemBioChem 12, 38–42 (2011).

38. Yi, L. et al. Engineering of TEV protease variants by yeast ER sequestration screening (YESS) of combinatorial libraries. Proc. Natl. Acad. Sci. 110, 7229–7234 (2013).

39. Piccolo, S., Dupont, S. & Cordenonsi, M. The biology of YAP/TAZ: Hippo signaling and beyond. Physiol. Rev. 94, 1287–1312 (2014).

40. Gera, N., Hussain, M. & Rao, B. M. Protein selection using yeast surface display. Methods 60, 15–26 (2013).

41. Benatuil, L., Perez, J. M., Belk, J. & Hsieh, C. An improved yeast transformation method for the generation of very large human antibody libraries. Protein Eng. Des. Sel. 23, 155–159 (2010).

42. Gera, N., Hussain, M., Wright, R. C. & Rao, B. M. Highly Stable Binding Proteins Derived from the Hyperthermophilic Sso7d Scaffold. J. Mol. Biol. 409, 601–616 (2011).

43. Isidro-Llobet, A., Álvarez, M. & Albericio, F. Amino Acid-Protecting Groups. Chem. Rev. 109, 2455–2504 (2009).

44. Chandra, K. et al. A Tandem In Situ Peptide Cyclization through Trifluoroacetic Acid Cleavage. Angew. Chemie - Int. Ed. 53, 9450–9455 (2014).

45. Kaiser, E., Colescott, R. L., Bossinger, C. D. & Cook, P. I. Color Test for the Detection of Free Terminal Amino Groups in the Solid-Phase Synthesis of Peptides. Anal. Biochem. 34, 595–598 (1970).

46. Menegatti, S. et al. Reversible Cyclic Peptide Libraries for the Discovery of Affinity Ligands. Anal. Chem. 85, 9229–9237 (2013).

47. Ramos-Tomillero, I., Rodriguez, H. & Albericio, F. Tetrahydropyranyl, a Nonaromatic Acid-Labile Cys Protecting Group for Fmoc Peptide Chemistry. Org. Lett. 17, 1680–1683 (2015).

48. Rachel, N. & Pelletier, J. Biotechnological Applications of Transglutaminases. Biomolecules 3, 870–888 (2013).

49. Schmidt, M., Toplak, A., Quaedflieg, P. J. L. M., Maarseveen, J. H. Van & Nuijens, T. Enzyme-catalyzed peptide cyclization. Drug Discov. Today Technol. 26, 11–16 (2017).

50. Deweid, L. et al. Directed Evolution of a Bond-Forming Enzyme: Ultrahigh-Throughput Screening of Microbial Transglutaminase Using Yeast Surface Display. Chem. - A Eur. J. 24, 15195–15200 (2018).

51. Sugimura, Y. et al. Screening for the Preferred Substrate Sequence of Transglutaminase Using a Phage-displayed Peptide Library. J. Biol. Chem. 281, 17699–17706 (2006).

52. Bacon, K., Burroughs, M., Blain, A., Menegatti, S. & Rao, B. M. Screening Yeast Display Libraries against Magnetized Yeast Cell Targets Enables Efficient Isolation of Membrane Protein Binders. ACS Comb. Sci. 21, 817–832 (2019).

53. Iglesias-Bexiga, M. et al. WW domains of the yes-kinase-associated-protein (YAP) transcriptional regulator behave as independent units with different binding preferences for PPxY motif-containing ligands. PLoS One 10, e0113828 (2015).

54. Yurimoto, H. et al. The Pro-peptide of Streptomyces mobaraensis Transglutaminase Functions in cis and in trans to Mediate Efficient Secretion of Active Enzyme from Methylotrophic Yeasts. Biosci. Biotechnol. Biochem. 68, 2058–2069 (2004).

55. Redding, K., Holcomb, C. & Fuller, R. S. Immunolocalization of Kex2 Protease Identifies a Putative Late Golgi Compartment in the Yeast Saccharomyces cerevisiae. J. Cell Biol. 113, 527–538 (1991).

